# Transposons and accessory genes drive adaptation in a clonally evolving fungal pathogen

**DOI:** 10.1101/2025.02.18.635021

**Authors:** Cristina López Díaz, Dilay Hazal Ayhan, Ana Rodríguez López, Lucía Gómez Gil, Li-Jun Ma, Antonio Di Pietro

## Abstract

Genomes of clonally reproducing fungal pathogens are often compartmentalized into conserved core and lineage-specific accessory regions (ARs), enriched in transposable elements (TEs). ARs and TEs are thought to promote pathogen adaptation, but direct experimental evidence is sparse. Using an evolve and re-sequence approach, we found that serial passaging of the cross-kingdom fungal pathogen *Fusarium oxysporum* through tomato plants or axenic media rapidly increased fitness under the selection condition. TE insertions were the predominant type of mutations in the evolved lines, with a single non-autonomous hAT-type TE accounting for 63% of total events detected. TEs inserted preferentially at sites of histone H3 lysine 27 trimethylation, a hallmark of ARs. Recurrent evolutionary trajectories during plate adaptation led to increased proliferation concomitant with reduced virulence. Unexpectedly, adaptive mutations in accessory genes strongly impacted core functions such as growth, development, quorum sensing or virulence. Thus, TEs and ARs drive rapid adaptation in this important fungal pathogen.

## Introduction

Fungal phytopathogens provoke devastating losses in agricultural productivity^1^, while opportunistic fungal infections in humans claim millions of lives each year^2^. Invasive fungal pathogens evolve predominantly as clones^3^, yet they can adapt rapidly to changing environments or new hosts^4,5^. The genomes of these pathogens are often compartmentalized into conserved core regions and lineage-specific accessory regions (ARs), which exhibit low gene density, high numbers of transposable elements (TEs), and can be horizontally transferred ^6–9^. In contrast to the core genome, which encodes essential housekeeping functions, ARs are often dispensable for life and encode adaptive functions such as virulence-related effectors or secondary metabolites^10^. First discovered in the pea pathogen *Nectria haematococca*^11^, ARs are found in a wide array of species including 5 out of the Top 10 fungal plant pathogens^12^ as well as animal and human pathogens^13,14^.

ARs and TEs have commonly been associated with high genetic plasticity and adaptation to ecological niches including plant hosts^3^. The term two-speed genome has been coined to describe the faster evolutionary rates of ARs versus core regions^15,16^. While comparative genomic surveys support a role of ARs and TEs in rapid adaptation to environmental stresses^17^ or plant hosts^18–20^, direct experimental proof for such a role is largely missing. Here we performed short term experimental evolution (STEE) to study adaptation in *Fusarium oxysporum* (*Fo*), a ubiquitous fungal pathogen whose genome is compartmentalized into well-defined core and ARs^21^. *Fo* displays a remarkable genetic and phenotypic plasticity in the absence of a known sexual cycle^22,23^, which is exemplified by the recent emergence of a highly aggressive clone named tropical race 4 (TR4) that threatens to wipe out the world’s most important staple crop, the Cavendish banana^24,25^. Collectively, *Fo* causes vascular wilts in more than 150 different crops and can also live as an endophyte on alternative host plants^26^. Its ecological versatility is further reflected by its capacity to provoke opportunistic infections in animal hosts, including humans^14^.

For STEE we chose the sequenced reference strain *Fo* f. sp. *lycopersici* 4287 (*Fol*4287), whose 60 Mb genome contains 19 Mb of ARs^21^. This single isolate can infect and kill tomato plants as well as immunodepressed mice and larvae of the insect host *Galleria mellonella*^27,28^. We found that serial passaging of *Fol*4287 through tomato plants or axenic media resulted in increased fitness in the selection condition, with TEs acting as the main drivers of genetic variation. Furthermore, TE insertions preferentially targeted ARs, causing adaptive mutations in accessory genes which strongly impacted fungal growth, development and virulence. Our results establish TEs and ARs as key drivers of rapid adaptation in this clonally evolving pathogen and reveal an unexpected role of ARs in the control of broadly conserved processes in fungi.

## Results

### Serial passaging of a clonal *Fo* isolate leads to rapid adaptation to the selection condition

Starting from a monosporic isolate of *Fol*4287 (thereafter called the ancestor clone), 10 serial passages were performed in different environmental conditions including tomato plants or plates with either complete (YPDA) or minimal medium (MMA) at 28°C or 34°C (Fig. 1a). The population bottleneck in all passaging conditions was around 1:20 (see Methods for description of *in planta* bottleneck calculation). Five independent replicate lines were performed for each condition. After the final passage, 10 monosporic isolates (called evolved isolates hereafter) were randomly picked from each evolved population and stored for phenotyping.

**Fig. 1.**
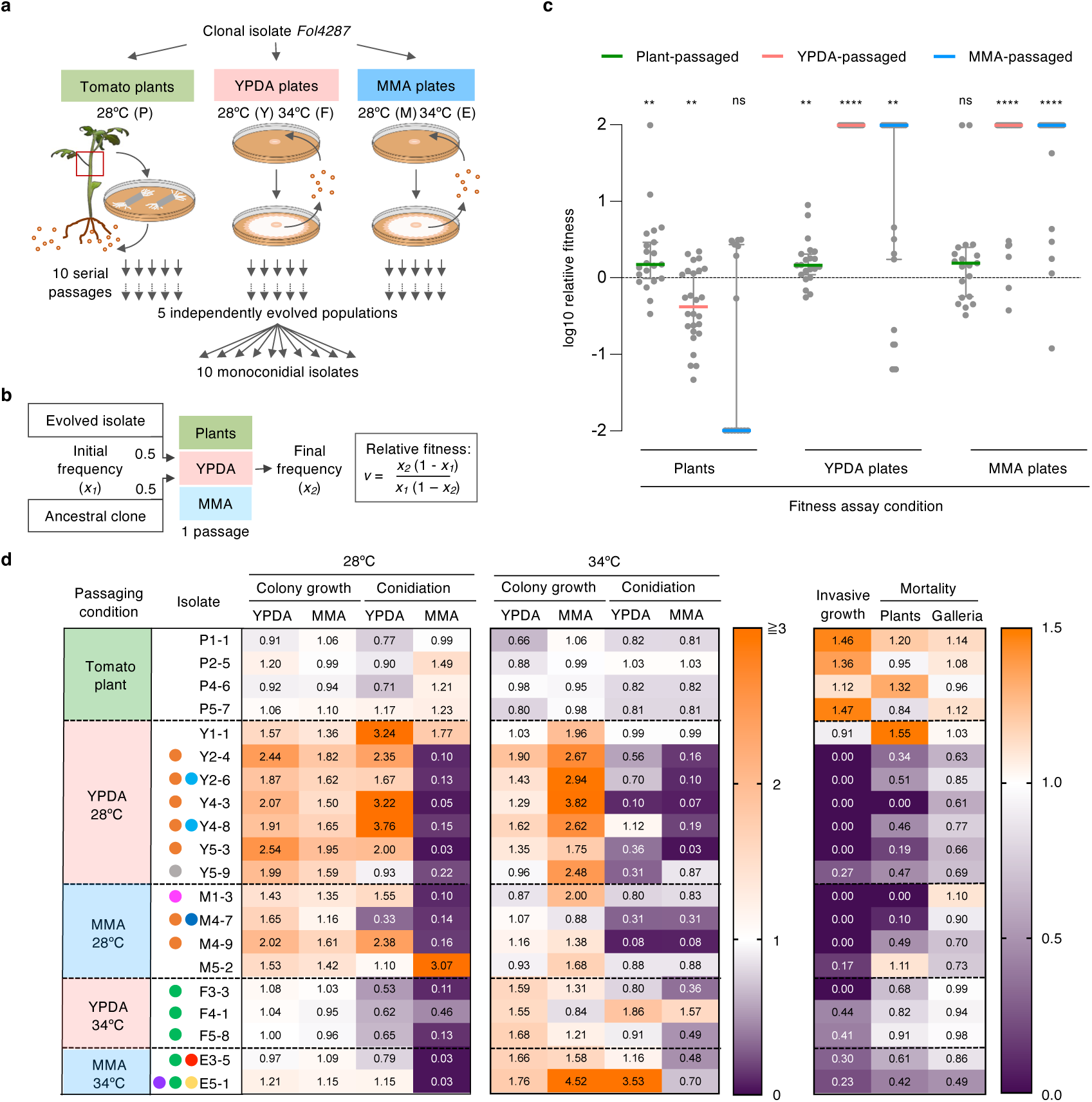
Fitness gains and trade-offs in clonally evolved populations of *Fusarium oxysporum*. **a.** Overview of the short term experimental evolution approach. *Fol4287* was submitted to 10 consecutive passages through tomato plants at 28°C; or through YPDA or MMA plates at 28°C or 34°C. For each condition, 5 independent replicate populations were performed. From each evolved population, 10 monoconidial isolates were obtained for fitness and phenotyping experiments. **b.** Schematic diagram of the pairwise competition assay used for fitness measurements. Relative fitness (*v*) was calculated as indicated. **c.** Repeated passaging leads to increased fitness in the selection condition. Scatter plots show log10 of mean relative fitness in the indicated conditions, of representative evolved isolates passaged through tomato plants (green, 4 isolates tested), YPDA (pink, 6 isolates) or MMA plates (blue, 4 isolates) at 28°C. Each dot is an experimental repeat with an evolved isolate. At least 4 independent experiments for each isolate were performed. Estimated detection limit for each isolate is 0.01. Colored lines are medians. Error bars, 95% confidence interval. **P < 0.01, ****P < 0.0001 according to one-sample Wilcoxon signed rank test, μ = 0. **d.** Heat maps showing values of the indicated phenotypes related to proliferation (left two panels) or invasion/pathogenicity (right panel) determined for the indicated evolved isolates, relative to the ancestral clone. Each row represents an evolved isolate, each column a phenotype. Colored dots next to isolates indicate adaptive mutations in recurrently selected genes (see Table 1 for legend).

We first asked whether repeated passaging through a constant environment leads to improved adaptation to the selection condition. To this aim, pairwise competition experiments were performed where evolved isolates were co-inoculated at equal frequency with the *Fol*4287 ancestor clone and submitted to a single passage under the same conditions used for STEE (Fig. 1b). Most evolved lines displayed significantly higher relative fitness under the selection condition than the ancestor clone (Fig. 1c). The fitness increase was particularly striking in the plate-evolved lines, many of which outcompeted the ancestor in a single plate passage. While most of the lines passaged through YPGA or MMA had increased relative fitness on both types of media, the selective advantage was stronger under the specific selection condition. Importantly, most plate-evolved lines exhibited reduced fitness in tomato plants compared to the ancestor clone (Fig. 1c), whereas plant-passaged lines did not show significant changes in fitness on plates. We conclude that STEE leads to rapid adaptation of *Fol*4287 to the selection condition, and that increased fitness on plates often entails a fitness loss on the host plant.

### Adaptation to plates results in increased proliferation at the cost of reduced invasion and virulence

To determine the phenotypic effects of adaptation, representative evolved isolates were scored for two traits associated with proliferation, colony growth speed and production of microconidia. Most plate-passaged isolates exhibited significantly increased colony growth on both YPDA and MMA, as well as increased conidiation on YPDA (Fig. 1d, Extended Data Fig. 1). Growth speed of most lines passaged at 28°C was increased both at 28°C and 34°C, while in lines passaged at 34°C it was only increased at 34°C. Unexpectedly, most plate-evolved isolates showed significantly reduced conidiation on MMA, including those selected on MMA (Fig. 1c, Extended Data Fig. 1).

Evolutionary adaptation to a given environment often entails trade-offs resulting in reduced fitness in a different condition^29^. Because most plate-passaged lines exhibited reduced fitness during colonization of the host plant tomato (Fig. 1c), the evolved isolates were scored for virulence-related traits such as invasive growth across cellophane membranes^30^ or mortality caused on tomato plants or the insect model host *Galleria mellonella*^31^. Strikingly, almost all plate-passaged isolates showed a marked decrease in these virulence-related traits (Fig. 1d, Extended Data Figs. 1 and 2). Two exceptions were the isolates derived from the YPDA-passaged line Y1 and the MMA-passaged line M5, both of which had slightly increased virulence on plants. Interestingly, these two isolates also differed from the other plate-evolved lines in exhibiting increased, rather than decreased conidiation on MMA (Fig. 1d). Plant-passaged isolates showed little or no changes in growth and conidiation on plates, but some exhibited a slight increase in invasive growth and virulence.

Pearson r correlation analysis of the phenotypes scored in the evolved isolates detected a positive correlation between colony growth speed on YPDA or MMA and conidiation on YPDA (Extended Data Fig. 3, box a). These proliferation-related phenotypes negatively correlated with virulence-associated traits such as cellophane invasion or mortality caused on tomato plants or Galleria, which in turn were positively correlated with each other (boxes b and c, respectively). This result demonstrates a trade-off between the selective advantages gained during plate adaptation and the fitness costs in virulence-related traits. Interestingly, conidiation on MMA correlated positively with virulence-related traits but negatively with plate-selected traits (boxes c and d), suggesting that the ability to sporulate under nutrient-poor conditions (MMA) could confer an advantage during infection of plant and animal hosts. Taken together, these results establish that adaptation of *Fol*4287 to growth on plates results in increased proliferation at the cost of reduced invasion and virulence on plant and animal hosts.

### TE insertions in ARs are the predominant type of mutation in experimentally evolved populations

To study the genetic changes underlying environmental adaptation, we generated whole genome sequences of 41 evolved populations obtained after passage 10, using Illumina paired-end reads. The genome of the *Fol*4287 ancestor clone (WT0) was also sequenced and assembled as the reference ^32^. More than 99% of the resequencing reads could be mapped to this reference genome, supporting the high quality of the sequence data (Supplementary Table 1). At a 10% coverage cut-off, a total of 320 *de novo* variants were detected across all passaged lines. More than 70% of the variants (228 events) correspond to TE insertion variations, while 23% are single nucleotide variations (74 events) and 5.5% are INDELs (18 events) (Fig. 2a panels A and B; Supplementary Tables 1 and 2). We independently confirmed 48 of the events (20 TIVs, 14 SNVs and 5 INDELs) by PCR, RFLP and/or Sanger sequencing. The number of TIVs in the populations passaged on YPDA and MMA plates at 34°C was 4.2 and 5.5 times higher, respectively, than in populations passaged on the same media at 28°C (p-values 0.003 and 0.051, respectively, according to paired Student’s t-test), suggesting that TIV frequency increases at high temperature (Fig. 2b). By contrast, the numbers of SNVs and INDELs were not affected by temperature. Among the 320 total mutations detected, 43% fell within coding, 13% within non-coding and 44% within intergenic regions (Extended Data Fig. 4a). The fraction of fixed or almost fixed mutations (allele frequency between 0.9 and 1) was significantly higher in plant-passaged than in plate-passaged populations (Extended Data Fig. 4b).

**Fig. 2.**
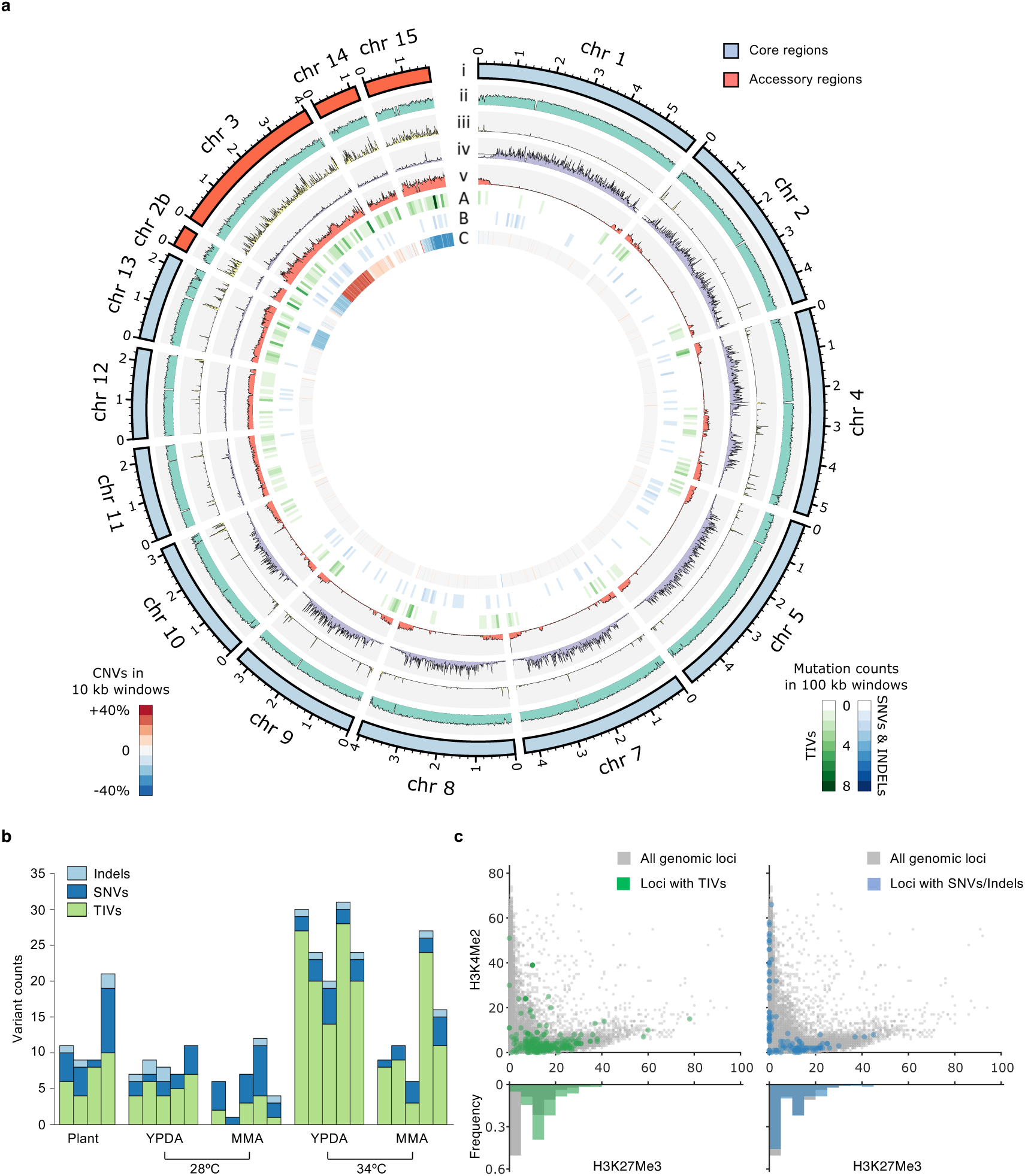
Transposon insertions in accessory regions are the predominant type of mutation in experimentally evolved populations. **a**. Circos plot summarizing the distribution of all the mutational events with allele frequency (AF) ≥ 0.1 detected in the passaged populations relative to the genomic region. (i) Chromosomes of the ancestral clone. Core genomic regions are in light blue, accessory regions in orange. (ii) GC % content; limits, 0.3-0.7. (iii) TE content; limits, 0-1. (iv,v) ChIP sequencing mapping of histone markers H3K4me2 associated with euchromatin (iv) or H3K27me3 associated with facultative heterochromatin (iv); limits, 0-50 (GEO accession: GSE121283)^33^. (A) Transposon insertion variants (TIVs; green) or (B) single nucleotide variants (SNVs)/Indels (blue) calculated by 100 kb windows. Color scale bars are in the lower right. (C) Averaged copy number variations (CNVs; blue, deletions; red, duplications) calculated by 10 kb windows. Color scale bar is in the lower left. **b.** Absolute counts of TIVs, SNVs and Indels (AF ≥ 0.1) in all the replicate populations passaged through the indicated conditions. **c.** Estimated histone marker levels of genomic loci with the indicated types of variants (AF ≥ 0.1) were calculated from ChIP-seq mapping data. Loci with TIVs or INDELs/SNVs are highlighted in green or blue, respectively. Upper panels, scatter plot of H3K4me2 and H3K27me3 read depths of 10 kb genomic windows. Lower panels, histograms of H3K27me3 read depth values.

In the compartmentalized genome of *Fol*4287^21^, the core regions correspond to the major parts of chromosomes 1 and 2 as well as to the entire chromosomes 4, 5, 7, 8, 9, 10, 11, 12 and 13, while the ARs encompass the entire chromosomes 3, 6, 14, 15, as well as minor sections of chromosomes 1 and 2 and several unmapped scaffolds^21^. We detected a significantly higher frequency of TIVs in ARs (4.1×10^-7^ TIVs/Mb/sample) than in core regions (1.3×10^-7^ TIVs/Mb/sample; p-value 0.041, paired student’s t-test), while SNPs or INDELs showed no such bias (Fig. 2a panel A). Furthermore, TIVs but not SNPs or INDELs were significantly enriched in regions of facultative heterochromatin carrying high levels of tri-methylation of lysine 27 on histone H3 (H3K27me3), a distinctive feature of ARs and subtelomeric regions^9,34^, but not in euchromatin regions marked with H3K4me2 (Fig. 2c; p-values 6.12×10^16^ and 0.28, respectively, according to one tailed Fisher’s Exact Test). We conclude that TIVs are the most frequent type of variants detected in the evolved lines and locate predominantly to ARs and subtelomeres.

### Evolved populations display recurrent copy number variations in ARs

Copy number variations (CNVs) represent a rapid mechanism for generating adaptive genetic variation in fungal pathogens^35,36^. Here we found that the evolved lines carried recurrent copy number variations (CNVs), including large segmental duplications and deletions. Almost all CNVs were associated with ARs or with the fast core chromosome 13 (Fig. 2a panel C) (P-value for non-random association between ARs with >10% CNV change in any sample 7.08×10^-253^ according to Fisher’s exact test). Three large-scale CNVs were detected in more than one evolved population and collectively encompass approximately half of the total ARs: 1) a 2.1 Mb duplication corresponding to part of chromosome 3 (detected in 9 of the 24 independently passaged lines); 2) loss of one copy of a duplicated region corresponding to chromosome 15 (in 14 lines); and 3) loss of one copy of a duplicated region corresponding to the small arm of chromosome 13 (in 13 lines). Recurrent CNVs were detected more frequently in plate-passaged than in plant-passaged populations. As a result of large-scale deletions, the average genome size of populations passaged through YPDA at 28°C, or through YPDA or MMA at 34°C, was reduced by 1.4, 5.9, or 1.5 Mb, respectively, compared to the ancestor clone, representing a loss of up to 10% of the genome (Extended Data Fig. 4c, Supplementary Table 1). By contrast, the populations passaged through plants or MMA plates at 28°C showed an average increase in genome size of 0.6 or 2.3 Mb, respectively, due to a large-scale duplication. Besides these recurrent CNVs, several smaller (< 500 kb) CNVs were detected in individual populations, either in core chromosomes (4, 9, 10) or accessory chromosomes (2, 3, 14).

To capture the dynamics of the recurrent CNVs, real time qPCR was performed across the serial passages of lines evolved at 28°C using specific primer pairs. The frequency of CNVs in the evolving population tended to increase rapidly, often leading to complete fixation (Extended Data Fig. 5). In summary, multiple CNVs, collectively encompassing approximately half of the ARs, were recurrently selected during experimental evolution indicating a possible role of structural dynamics of ARs in environmental adaptation of *Fol*4287.

### Almost 80% of TIVs are caused by a single miniature DNA transposable element

Among the total 228 TIVs detected in the passaged populations, close to 90% (203 events) were caused by only two DNA TEs: *Hormin* (181 events; 79.4%) and *MarCry-1* (22 events, 9.6%) (Fig. 3a). The most active TE *Hormin* is a non-autonomous miniature terminal inverted repeat (TIR) element (MITE) of 759 bp length, derives from the full-length hAT superfamily transposon *Hornet* ^37,38^. The second most active TE, *MarCry-1_FO*, is an autonomous TIR element from the TcMariner superfamily^37,39^. Interestingly, *Hormin* and *MarCry-1_FO* are also the TIR elements with the highest copy number in the *Fol*4287 genome (77 and 46 copies, respectively)^40^, suggesting a possible correlation between mobility and the number of TE copies in the genome.

**Fig. 3.**
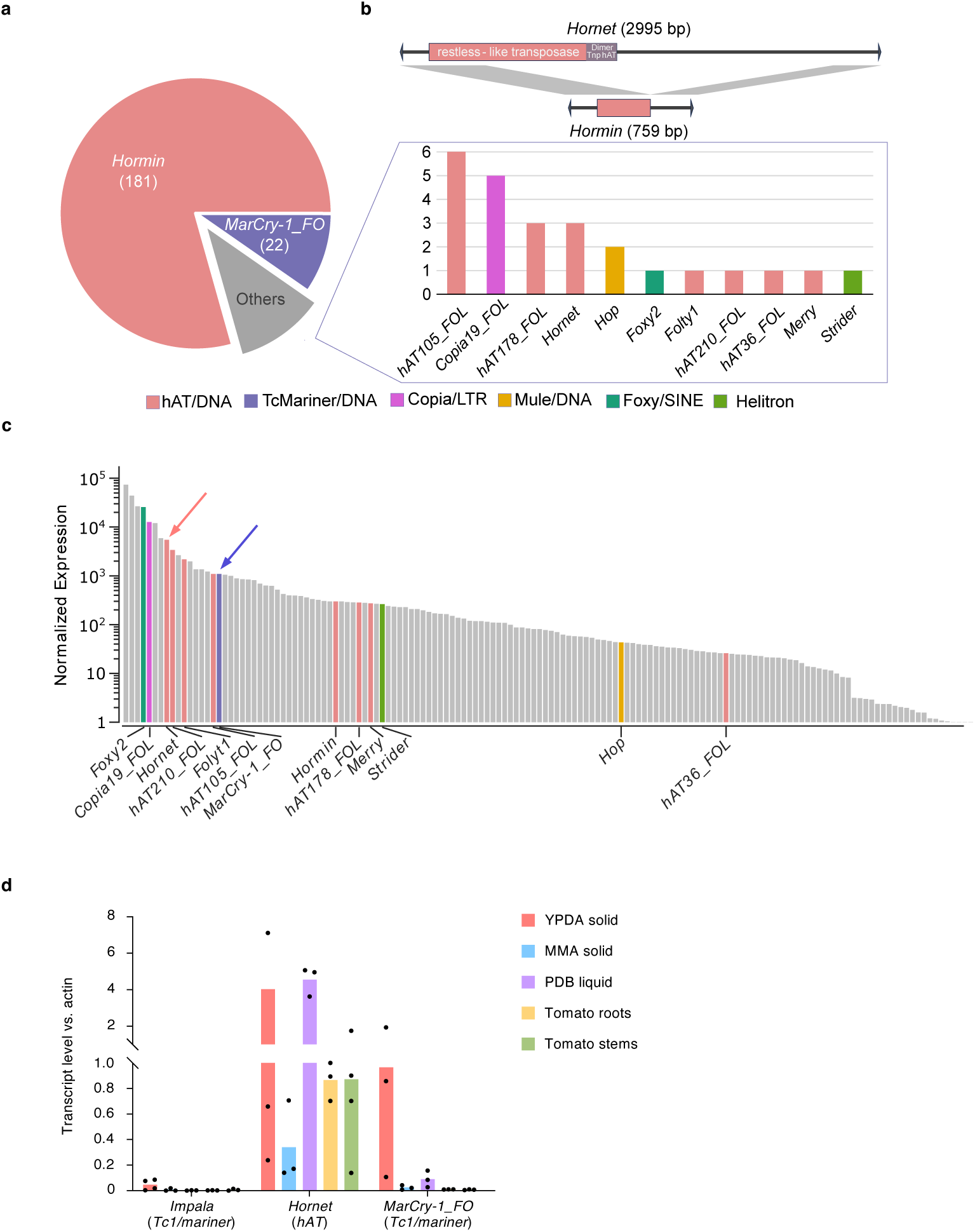
Almost 80% of TIVs detected in evolved populations are caused by a single non-autonomous MITE. **a.** Proportion and total numbers of TIVs (AF ≥ 0.1) caused by the indicated transposable elements (TEs), detected in the evolved populations. Color code corresponds to indicated TE superfamilies. **b.** Schematic view of the full-length *hAT-*type TE *Hornet* and its non-autonomous miniature version *Hormin*. Homologous regions are in grey. The restless-like transposase and the *hAT* family C-terminal dimerization regions (Dimer_Tnp_hAT) are indicated. Arrowheads represent the 16 bp terminal inverted repeats (TIRs). **c.** Normalized expression of different TEs during growth of *Fol4287* in liquid YPD medium was measured by RNA-seq. Active TEs detected as TIVs in experimentally evolved lines are colored according to superfamily (see a) Arrows point to the TEs *Hornet* and *MarCry-1_FO*. **d.** Relative transcript levels of the indicated transposases, measured by RT-qPCR of cDNA obtained from *Fol4287* grown on solid YPDA or MMA, or in liquid PDB medium; or from roots or stems of infected tomato plants. Levels are normalized to those of the *Fol4287* actin gene. Dots are independent biological repeats.

DNA TEs mobilize through the activity of transposases, which catalyze DNA cleavage and TE insertion through a “cut and paste” process^41^. Non-autonomous MITEs such as *Hormin* lack a functional transposase and thus depend on the enzyme encoded by the corresponding full-length element^42^. *Hormin* is likely mobilized by the restless-type transposase of its full-length counterpart *Hornet* (Fig. 3b), which is present in 24 copies in the *Fol*4287 genome^40^. Interestingly, *Hornet* was the most highly expressed DNA TE and ranked among the most highly expressed TEs overall (Fig. 3c). Transcript levels of *Hornet* transposase were higher than those of the highly expressed actin gene during growth of *Fol*4287 on solid or liquid media, or during infection of tomato plants (Fig. 3d). The second most active TE, *MarCry-1_FO,* also ranked among the most highly expressed DNA TEs, although its transcript levels were approximately one order of magnitude lower than those of *Hornet* (Fig. 3c,d). Notably, retrotransposons such as LTRs, LINEs, SINEs, which ranked among the TEs with the highest expression levels and copy numbers in the *Fol*4287 genome (Fig. 3c)^40^, only accounted for 3% of the TIVs detected in the evolved populations (Fig. 3a).

### Adaptation to plates follows recurrent evolutionary trajectories

The finding that most evolved lines displayed increased fitness under the passaging conditions (Fig. 1c) suggested the presence of strong positive selection. In line with this, 44.6% of the SNPs detected in evolved lines had consequences on the protein sequence. Among the SNPs located in coding sequences (CDSs), 60.6% were missense, 18.2% nonsense and only 21.2% were synonymous mutations (Supplementary Table 2). To assess the repeatability of evolution under selection and to identify the mutations that drive adaptation, we searched for genes mutated more often than expected by chance. If the 320 mutations were distributed randomly over the 21,560 predicted genes of *Fol*4287, 2.34 genes are expected to be hit more than once in independent lines. Instead, we found that 14 genes were hit two or more times, representing a total of 43 independent mutational events (called recurrent mutations thereafter; Table 1). Furthermore, mutations in these genes had a significantly higher fixation rate (11/43, 25.6%) compared to all other nonsynonymous mutations (40/277, 14.4%, P-value 0.0738, Fisher exact test). Eight of 14 genes are located on core regions (20 events), while 6 are on ARs (23 events). Strikingly, all recurrent mutations in AR genes are TIVs, while only half of the recurrent mutations in core genes are TIVs (Table 1). More than half of the recurrent mutations (24 events in 7 genes) likely result in loss of function.

**Table 1.**
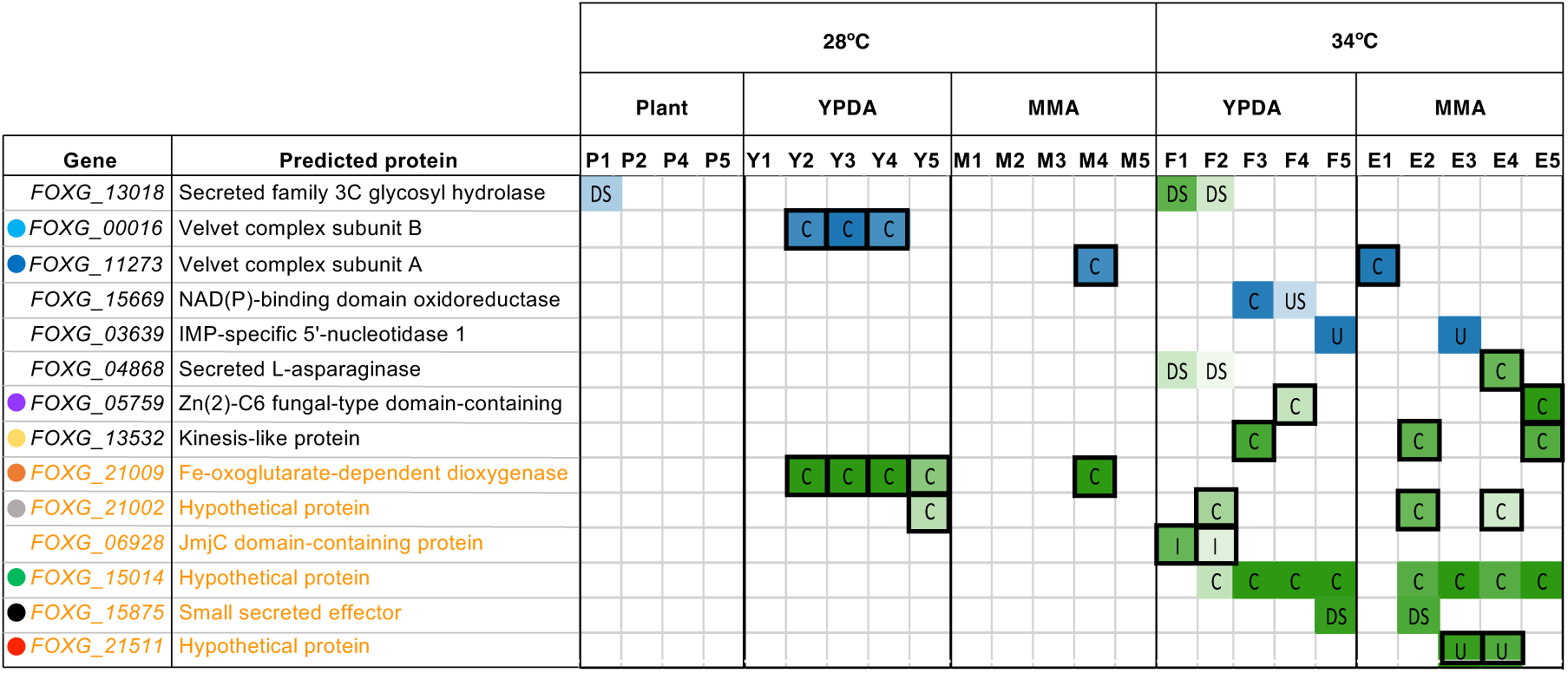
Core and accessory genes recurrently mutated in independently evolved populations. List of genes mutated in more than one passaged population (AF ≥ 0.1). Columns represent independent populations passaged through the indicated condition. Genes in orange are located on accessory regions. Green and blue heatmaps represent allele frequencies of TIVs and SNVs/INDELs, respectively. Letters in cells represent the position of the mutation relative to the gene. C: coding, U: UTR, I: Intron, DS: downstream, US: Upstream. Boxed in black are mutations likely selected for loss of function (TIVs in CDSs, UTRs or introns; Indels or nonsense mutations in CDSs). Functional predictions were obtained by blastp and/or alphafold/foldseek searches. Colored dots next to genes indicate recurrent mutations detected in evolved isolates used for phenotyping (see Fig. 1d).

Importantly, all except one of the recurrent mutations were detected in plate-passaged lines, where they account for > 15% of the total events, strongly suggesting that adaptation of *Fol*4287 to the plate environment follows repeated evolutionary trajectories. The most frequent trajectory was a defined sequence of successive loss-of-function mutations in two genes, *FOXG_21009* and *velB* (*FOXG_00016*) (Fig. 4a). Four of the 5 lines passaged on YPDA acquired a *Hormin* insertion in the CDS of *FOXG_21009* and 3 of these subsequently acquired a SNV or INDEL in the *velB* CDS resulting in a premature stop codon (Fig 4b). *FOXG_21009* encoding a hypothetical protein is on accessory chromosome 3, while *velB* is on core chromosome 1 and encodes the subunit B of the conserved Velvet complex, a master regulator of fungal development and secondary metabolism^43^. Interestingly, 2 of the 5 lines passaged on MMA plates also carried a TIV in *FOXG_21009* and one of them additionally acquired a nonsense mutation in the *veA* gene (*FOXG_11273*), which encodes the subunit A of the Velvet complex (Extended Data Fig. 6a). The fact that all plate-passaged populations carrying a fixed TIV in *FOXG_21009* subsequently acquired a secondary mutation in *velB* or *veA* (Fig. 4b and Supplementary Table 2, events T091, T092, S002, S004) strongly suggests an epistatic relationship between these genes.

**Fig. 4.**
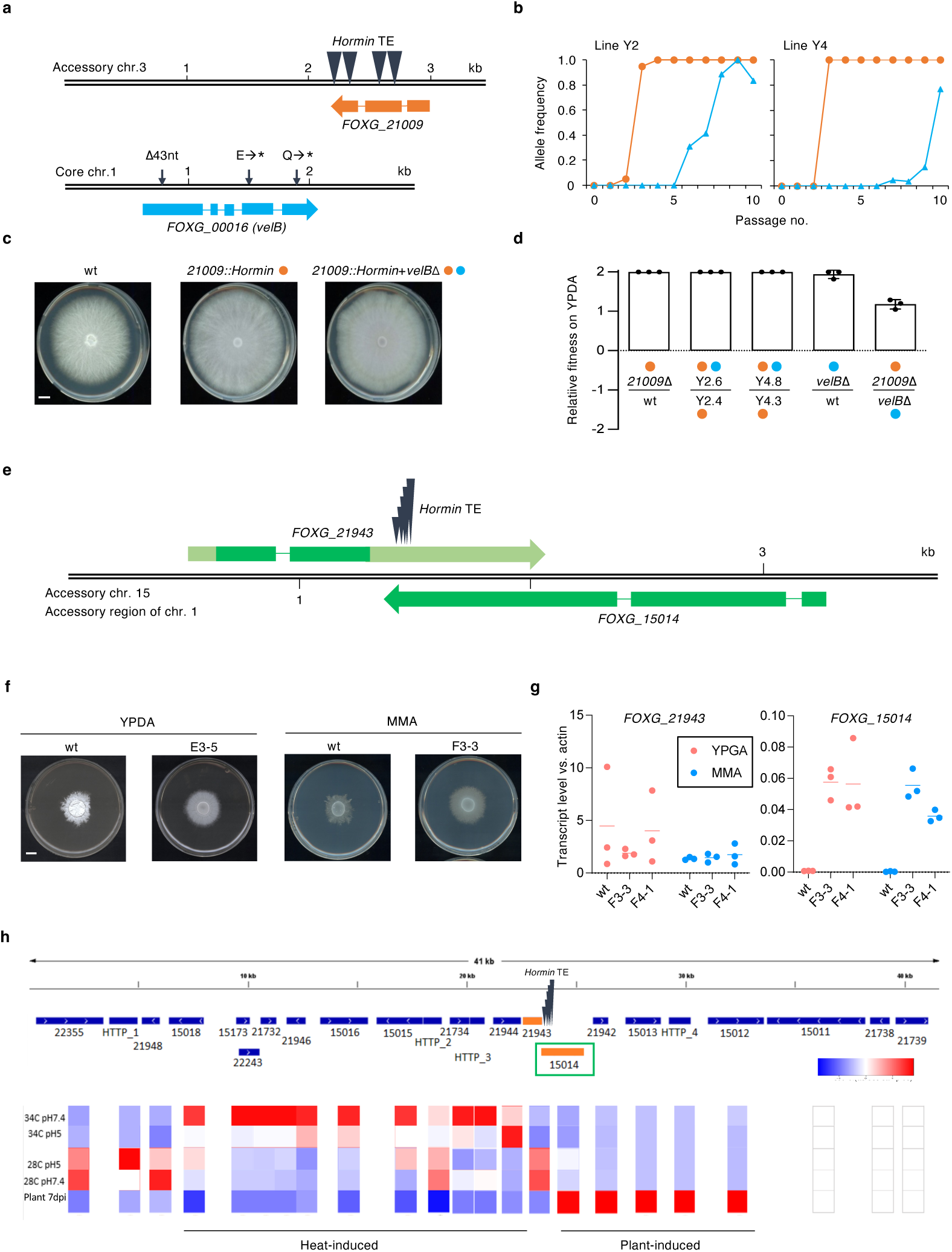
Adaptive mutations in accessory genes selected during serial passages on plates. **a-d.** Successive adaptive mutations selected in multiple independently evolved populations passaged on YPDA plates at 28°C. **a.** (Upper) *Hormin* insertions in four independent lines map to the CDS of the *FOXG_21009* gene. (Lower) Indels or SNVs in three independent lines resulting in premature stop codons in the *velB* gene. **b.** Allele frequency dynamics of mutations in *FOXG_21009* (orange) and *velB* (blue) in two independent YPDA-passaged populations (Y2 and Y4). **c.** Colony phenotype on YPDA plates of the ancestor (wt) and YPDA-passaged isolates carrying a *Hormin* insertion in *FOXG_21009* (orange dot), either alone or in combination with a mutation in *velB* (blue dot). Scale bar, 1 cm. **d.** Adaptive mutations incrementally increase fitness on plates. Bars show log10 of the mean relative fitness in pairwise competition assays on YPDA between the indicated isolates derived from the two independently evolved populations Y2 and Y4; wt is the ancestor; Y2.4, Y2.6, Y4.3 and Y4.8 are evolved isolates; *21009*Δ and *velB*Δ are targeted deletion mutants. Orange and blue dots next to isolates indicate mutations in *FOXG_21009* and/or *velB*, respectively. Estimated detection limit for each isolate is 0.01. Black dots are independent experimental repeats. Error bars, SEM. **e.** *Hormin* insertions in independent populations passaged at 34°C on YPDA or MMA map to a small region in the genes *FOXG_21943* (3’-UTR) and *FOXG_15014* (3’-end of CDS). **f.** Colonies of the ancestor (wt) and the indicated evolved isolates carrying a TIV depicted in (e) were grown on the indicated media at 34°C. **g.** *Hormin* insertion in the 3’-end of the *FOXG_15014* CDS results in its transcriptional de-repression. Relative transcript levels of *FOXG_21943* and *FOXG_15014* in the indicated strains grown on YPDA or MMA plates at 34°C, determined by RT-qPCR and normalized to those of the actin gene. Dots are independent biological repeats. Colored lines are medians. **h.** Recurrently selected *Hormin* insertions map to the junction of two differentially regulated gene blocks, one induced at 34°C (*FOXG_15018* through *FOXG_21944*) and the other during plant infection (*FOXG_15014* through *FOXG_ 15012*). Heat map shows relative transcript levels of the genes located in the depicted 41 kb region, measured by RNA-seq during growth of *Fol*4287 in liquid YPD at 34°C or 28°C at the indicated pH; or in tomato roots at 7 days post inoculation (dpi). Green box indicates the *FOXG_15014* gene carrying recurrent *Hormin* insertions.

Both mutations were associated with marked changes in colony morphology (Fig. 4c, Extended Data Fig. 6b). While TIVs in *FOXG_21009* resulted in increased radial growth, the subsequent mutations in *velB* or *veA* caused a flat colony phenotype with reduced aerial mycelium, previously described for *Fol*4287 *velvet* mutants^44^. To investigate the loss-of-function mutations in these two genes, we independently created null mutants in the ancestor *Fol*4287 (Extended Data Fig. 7). The *21009* and the *velB*Δ mutants efficiently outcompeted the ancestor clone on YPDA plates (Fig. 4d), confirming that loss of function of these genes increases relative fitness of *Fol*4287 on plates. Importantly, isolates from two independently evolved lines carrying mutations in both genes (Y2-6 and Y4-8) fully outcompeted isolates from the same lines carrying only the *FOXG_21009* mutation (Y2-4 and Y4-3, respectively), thereby confirming that the mutation in *velB* further increases fitness of the *FOXG_21009* mutant on YPDA (Fig. 4d). Together with the finding that *21009*Δ successfully outcompeted *velB*Δ (Fig. 4d), these results explain the defined order in which these two recurrent mutations were selected during plate adaptation.

Among the 10 independent lines passaged at 34°C, 8 lines carried a *Hormin* insertion in a small 51 bp region located at the 3’-overlap of two genes: *FOXG_15014* encoding a hypothetical protein and *FOXG_21943* encoding a putative prolyl hydroxylase (Fig. 4e, Table 1). The two genes are located on an AR present in two copies in the *Fol*4287 genome, one on chromosome 15 and one on the accessory part of chromosome 1. Interestingly, the TIV is in heterozygosis in some of the evolved lines and in hemizygosis in others (due to loss of chromosome 15), suggesting that the mutation is dominant. In line with this, all evolved lines carrying this TIV exhibited increased colony growth at 34°C on YPDA and MMA, regardless of whether the mutation was present in hetero- or hemizygosis (Fig. 4f) (Extended Data Fig. 6f). Consistent with a gain of function, transcript levels of the *FOXG_15014* gene in two independent isolates carrying the TIV were up to 135-fold higher than in the ancestral clone during growth on YPGA or MMA plates (Fig. 4g). We further noted that the 51 bp region carrying the independent TIVs is located at the junction of two differentially regulated gene blocks, a block of 9 genes induced by high temperature (34°C) and another block of 5 genes induced during tomato plant infection (Fig. 4h).

The MMA-selected line M1 carried a missense mutation (G183S) in the *fadA* gene (*FOXG_09359*), which became rapidly fixed in the population (Extended Data Fig. 6c, d). The *fadA* ortholog in *Aspergillus nidulans* encodes an alpha-1 subunit of a guanine nucleotide-binding protein (G alpha) that regulates vegetative hyphal growth^45^. The G183S substitution prevents interaction of G alpha-1 with its cognate regulator of G protein signaling, resulting in constitutive activation of signaling and increased vegetative growth^46,47^. We found that line M1 displayed strong hyphal proliferation and a characteristic fluffy colony phenotype (Extended Data Fig. 6e) resembling that reported in the *A. nidulans* G183S mutant^45^. Interestingly, recurrent selection of the G183S mutation in FadA was previously observed in the ascomycete *Podospora anserina* upon serial passaging through submerged culture^48^. Importantly, most evolved lines carrying recurrent plate-selected mutations displayed a significant reduction in virulence-related traits such as invasive growth on cellophane or apple slices and caused significantly less mortality on tomato plants or *Galleria mellonella* (Fig. 1d, Extended Data Figs. 1-3). Together, these results demonstrate that adaptation of *Fol*4287 to plates follows reproducible evolutionary trajectories which result in increased proliferation in the plate environment but reduced virulence on plant and animal hosts.

### A previously unknown accessory gene controls conserved functions in fungal development and virulence

Almost half of the adaptive mutations selected on plates are in genes located on ARs. This result was unexpected, not only because ARs have a lower gene density than core regions, but also because accessory genes are thought to function mainly in adaptation to specific niches such as the plant host^21,49^. Here we focused on the recurrently mutated accessory gene *FOXG_21009*, whose predicted product has homologs in different *Fo* isolates as well as in other *Fusarium* species and ascomycetes such as *Verticillium, Colletotrichum, Penicillium* or *Metarhizium* (Fig. 5a). Structure prediction with Alphafold, followed by a database search with the structural alignment tool Foldseek^50^, identified a Fe(II)- and 2-oxoglutarate (2OG)-dependent dioxygenase domain-containing protein from *Pseudomonas aeruginosa*^51^ as the closest hit with a known function (Fig. 5b). This class of enzymes are found in all kingdoms of life and typically catalyze the oxidation of an organic substrate using a dioxygen molecule, as well as ferrous iron as a cofactor and 2OG as a co-substrate^52^.

**Figure 5.**
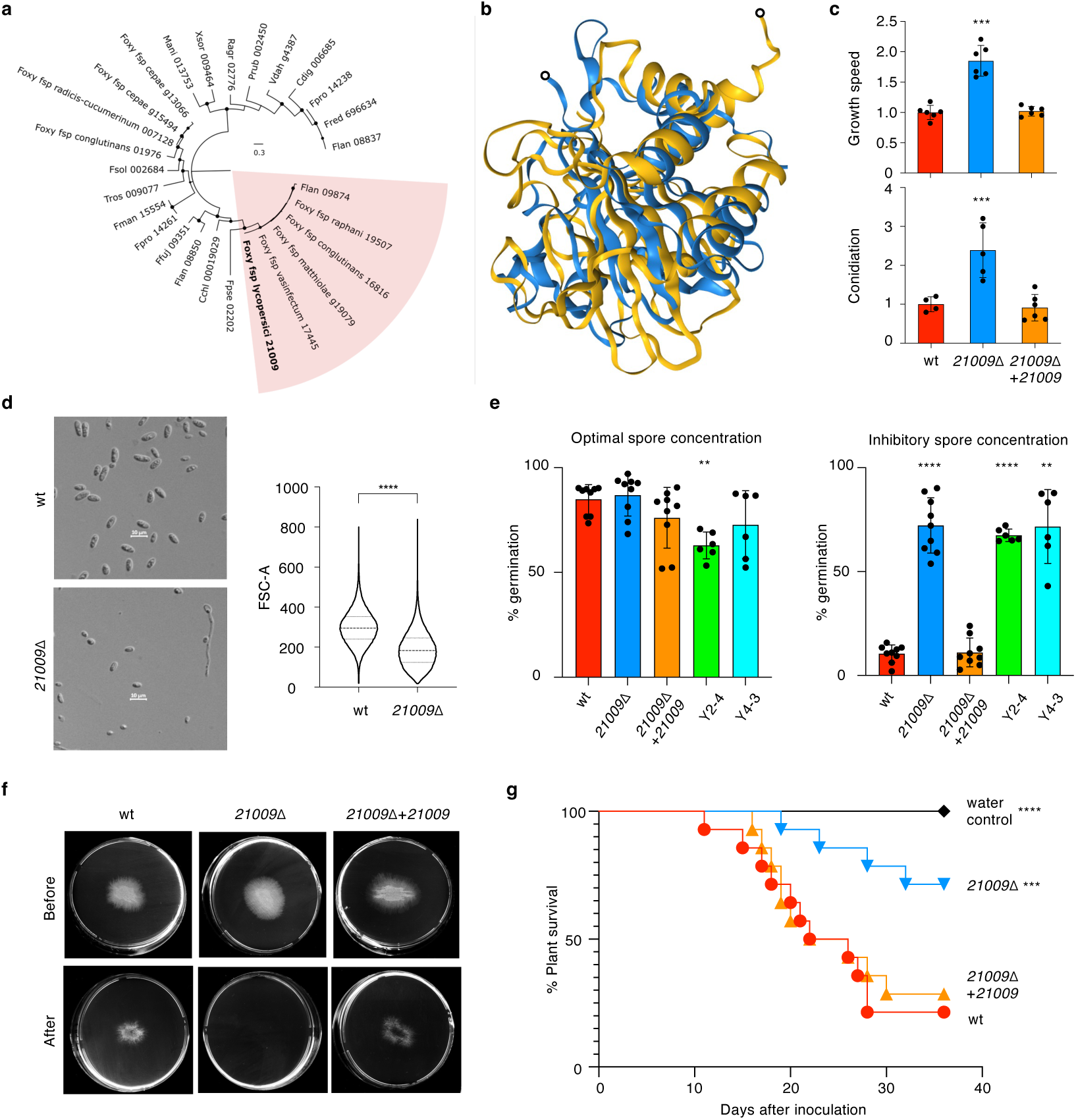
Accessory gene *FOXG_21009* encodes a predicted Fe(II) 2OG dioxygenase regulating fungal growth, development, quorum sensing and virulence. **a.** Maximum likelihood phylogenetic tree based on the aligned amino acid sequence of predicted FOXG_21009 protein homologs in the indicated fungal species. *Fo* clade is highlighted in pink. Numbers indicate NCBI IDs. Abbreviations of species names are listed in Supplemental Table S1. **b.** Structure of the FOXG_21009 protein (blue) predicted by Alphafold (amino acids 4 to 208), superimposed with the Fe(II) 2OG dioxygenase domain-containing protein PA1894 of *Pseudomonas aeruginosa* (yellow) using Foldseek. TM score 0.397, RMSD 6.02. N-termini were removed from the model (black circles). **c.** Growth speed and conidiation of the wild type (wt), the *21009*Δ knockout mutant and the complemented strain on YPDA plates at 28°C, normalized against the mean of wt. Error bars, SD. ***P < 0.001 versus wt according to one-way ANOVA. **d.** (Left) Representative micrographs of microconidia of the indicated strains. Scale bar, 20 μm. (Right) Violin plot of forward scatter linear-scaled values of microconidia from the indicated strains (right). Dashed line represents median, dotted lines quantile at 25% and 75%. n = 10^5^ conidia ****P < 0.0001 according to Mann-Whitney test. **e.** Percentage germination of microconidia of the indicated strains at optimal (left) or inhibitory spore concentration (right). n = 200 conidia **P < 0.01, ****P < 0.0001 versus wt according to one-way ANOVA. **f.** Invasive growth of the indicated strains through cellophane membranes placed on PDA plates. Plates were imaged after 2 days (Before), then the cellophane with the fungal colony was removed and plates were incubated for an additional day to visualize mycelium penetrated through cellophane (After). Scale bar, 1 cm. **g.** Kaplan-Meier plots showing survival of tomato plants inoculated with the indicated strains. n = 20 plants. ***P < 0.001, ****P < 0.0001 versus plants inoculated with wt according to Log-Rank test.

Targeted deletion of *FOXG_21009* in *Fol*4287 resulted a two-fold increase in colony growth speed and conidiation on YPDA compared to the wild type and the complemented strain *21009Δ*+*21009* (Fig 5c), mimicking the phenotype of the plate-evolved lines carrying a TIV in *FOXG*_*21009*. We noted that microconidia of *21009Δ* were significantly smaller than those of the wild type (Fig. 5d) and could germinated at high concentrations (8.6 x 10^7^ conidia/ml) that are inhibitory for the wild type and the complemented strain (Fig. 5e). This suggests that loss of *FOXG*_*21009* affects conidial quorum sensing, a self-sensing process previously reported in *Fol*4287^53^. While the frequency of hyphal fusion of *21009Δ* was similar to the wild type and the complemented strain (Supplementary Table 1), the mutant was severely impaired in invasive growth and exhibited significantly reduced virulence on tomato plants and Galleria *larvae* (Fig. 5 g,h, Extended Data Fig. 2). Thus, the previously uncharacterized accessory gene *FOXG_21009* regulates conserved fungal processes such as growth, development and virulence.

## Discussion

### TEs are major drivers of environmental adaption in *F. oxysporum*

Asexually evolving fungal pathogens with compartmentalized genomes display remarkable adaptive plasticity, but the underlying mechanisms have rarely been addressed experimentally^4,5^. Here we confirmed the capacity of a clonally propagating *Fo* isolate to rapidly adapt to new environments, significantly increasing its fitness after only 10 serial passages through the selection condition. TIVs accounted for more than 70% of the mutations found in the passaged populations, both on axenic media and in tomato plants. These results provide experimental proof of TEs acting as the main drivers of adaptive evolution in a fungus with a compartmentalized genome. Previous evidence for such a role was based largely on comparative genome analyses of two-speed genome fungal pathogens such as *V. dahliae* and *Z. tritici* ^18,20,54^. Strikingly, almost 80% of the TIVs corresponded to insertions of a single non-autonomous MITE, which caused 63% of the total mutations detected in the evolved lines. MITEs can achieve very high mobility rates by sequestering the transposition machinery encoded by the autonomous TE, allowing them to successfully outcompete the full-length element^42,55^. This could explain our finding that the full-length TE *Hornet* only caused 3 TIVs in the passaged lines, whereas the corresponding MITE *Hormin* caused 181 TIVs. *Hornet* was the most highly expressed DNA TE in *Fol*4287, suggesting that high levels of transposase produced by the full-length element can lead to massive mobility rates of the corresponding MITE. How each of the 24 *Hornet* copies contributes quantitatively to total transposase levels remains subject to further studies.

TIV frequencies were significantly higher in ARs than in core regions. Moreover, TIVs were enriched in facultative heterochromatin regions marked with H3K27me3, a distinctive feature of ARs and subtelomeres^9,34^. Increased TIV frequencies were also detected in the three smallest core chromosomes, also known as fast core chromosomes, which carry higher levels of H3K27me3 than the remaining core chromosomes^33^. These results suggest that TE insertions preferentially target ARs, where they represent the driving force of adaptation in the compartmentalized *Fol*4287 genome. The reason for the increased TIV rate associated with ARs or H3K27me3 is unknown. Previous studies in animals and plants detected TE insertion preferences for regions enriched in specific chromatin states, including H3K27me3^56–58^. Whether H3K27me3 is directly involved in targeting TEs such as *Hormin,* or whether TE insertion into H3K27me3-rich regions is facilitated by other factors associated to this epigenetic mark remains to be determined.

Regardless of the underlying mechanism, our results provide an explanation for the frequently observed enrichment of TEs in ARs. TE insertions have been associated with rapid sequence diversification and epigenetic regulation of virulence-related accessory genes such as effectors^16,19,21^, as well as increased tolerance to environmental stressors^17^. The high TE insertion rate detected in our study suggests a possible mechanism for the close physical association of TEs and accessory genes, including the enrichment of MITES such as *Hormin* or *Miniature Impala* (*mimp*) in the CDSs or promoters of effector genes reported in *Fol* and in other fungal pathogens ^19,38,59^. An intriguing observation was the selection of independent TIVs at the 3’-end of the *in planta*-induced gene *FOXG_15014* in 8 of the 10 lines passaged at 34°C, which caused a marked increase in gene transcript levels during growth on plates at 34°C. This finding, together with the finding that these TIVs map to the junction of two differentially regulated gene blocks, further supports a link between chromatin landscape, TE insertion and transcriptional regulation of adjacent accessory genes.

Importantly, the TIV frequency in the lines passaged at 34°C was significantly higher than in those passaged at 28°C, suggesting that high temperature promotes TE mobility in *Fol*4287. While the mechanism of temperature-dependent TE mobility is currently unknown, our findings are in line with previous studies showing increased TE mobility at high temperature in the human fungal pathogen *Cryptococcus deneoformans*^60,61^. These results suggest that the crucial role of TEs in environmental adaptation of fungal pathogens could become even more prominent under conditions of high temperature, such as those encountered during infection of mammalian hosts or in the context of global warming.

### Experimental evolution identifies novel accessory genes functioning in conserved fungal processes

Experimental evolution with multiple independent lineages often results in the selection of adaptive mutations in the same genomic loci^62,63^. We found independent *Hormin* insertions in the CDSs of *FOXG_21009* in 4 out of 5 lineages passaged on YPDA at 28°C, and of *FOXG_15014* in 8 out of 10 lineages passaged at 34°C. Furthermore, we detected evidence for epistasis between mutations in *FOXG_21009* and in the *velB* or *veA* gene encoding different subunits of the Velvet complex, suggesting that the benefit conferred by the second mutation depends on the genetic interaction with the previous mutation^64^. The dramatic phenotypic changes observed in plate-passaged lines of *Fol*4287 are in line with historic studies reporting highly variable morphologies arising during plate culture of different *Fo* strains, as well as repeated trajectories leading from colonies with profuse aerial mycelium and high virulence to variants with flat mycelial growth and reduced virulence^23^. Together, these results suggest that mutations favouring colony proliferation on plates entail important fitness trade-offs in virulence-related functions.

Several recurrent mutations selected on plates caused a loss of gene function, including the TIVs in *FOXG_21009* and the SNPs/indels in the *velvet* genes. Loss-of-function mutations have been proposed as a rapid evolutionary response during adaptation to new environments^65^. For example, deletion mutants in 630 non-essential genes of *S. cerevisiae* showed increased fitness in at least one of six different environments tested, although only 20 of these were fitter in all six environments^66^. Our results suggest that certain genes, which make *Fo* a successful generalist capable of thriving in a wide range of environments, could limit its fitness in specific niches such as those encountered during growth on plates.

About half of the adaptive mutations selected during plate adaptation target genes located on ARs. Accessory genes were previously thought to function mainly in host-pathogen interactions, such as host-specific virulence effectors or secondary metabolite gene clusters^21,67^. Our results reveal a previously unrecognized role of accessory genes in conserved fungal processes such as vegetative growth and development. One example is *FOXG_21009*, which encodes a putative Fe(II) 2OG dioxygenase. This class of proteins are widespread in bacteria and eukaryotes, where they function in a variety of processes ranging from hypoxia in metazoans^68^ to plant hormone metabolism^69^ or antibiotic biosynthesis in fungi^70^. While the precise biochemical function of the FOXG_21009 protein remains to be elucidated, phenotyping of the deletion mutant revealed a key role of this accessory gene in core functions such as hyphal growth, sporulation and conidial quorum sensing, as well as in pathogenicity-related traits such as invasive hyphal growth or killing of plant and animal hosts. We speculate that ARs could harbour additional, so far uncharacterized genes functioning in broadly conserved fungal processes, representing a largely untapped source for discovery of new antifungal targets. In summary, our passage and re-sequence experiments identify TEs and ARs as the main evolutionary force driving environmental adaptation in a fungal pathogen with a compartmentalized genome. This knowledge could be useful for designing new strategies to tackle fungal diseases in the context of global warming, environmental stresses and host transitions, both in agricultural and clinical settings.

## Methods

### Fungal strains, culture conditions and targeted gene knockout

*Fusarium oxysporum* f. sp. *lycopersici* race 2 isolate 4287 (*Fol*4287; FGSC 9935) was used as parental strain in all experiments. Strains were stored as microconidial suspensions in 30% glycerol at −80°C. To obtain fresh microconidia or mycelia for DNA extraction, fungal strains were grown in potato dextrose broth (PDB) at 28°C with orbital shaking at 170 rpm as described^1^. Targeted gene replacement through protoplast transformation with the hygromycin B resistance cassette (Hyg^R^), identification of knockout mutants by PCR and Southern blot analysis, and complementation by co-transformation with the wild type *FOXG_21009* allele and the phleomycin resistance cassette (Phleo^R^) were done as described^1^.

### Experimental evolution by serial passages through tomato plants or plates with different growth media

A clonal isolate of *Fol*4287 obtained by monoconidial isolation (hereafter called the ancestor clone or wt) was submitted to 10 consecutive passages through tomato plants, or on plates containing either yeast extract peptone dextrose agar (YPDA; 0.3 % w/v yeast extract, 1% w/v bactopeptone, 2% w/v glucose, 1.5% w/v agar, pH 5.5) or minimal medium agar (MMA; 0.1% w/v KH_2_PO_4_, 0.05% w/v MgSO_4_·7H_2_O, 0.05% w/v KCl, 0.2% w/v NaNO_3_, 3% w/v sucrose, 1,5% w/v agar, pH 5.5). For *in planta* passaging, roots of 2-week-old tomato plants (*Solanum lycopersicum* cultivar Monika) were dip-inoculated as described^1^, by submerging them 30 min in a suspension containing 5 x 10^6^ microconidia/ml, planted in minipots containing vermiculite, and maintained in a growth chamber at 28°C and a photoperiode of 14 h light/10 h dark. After 14 days the stems were surface sterilized during 2 min in 1% (w/v) NaOCl and rinsed twice for 2 minutes in sterile distilled water. Three stem pieces of approximately 1 cm length were cut with a sterile blade, placed on Potato Dextrose Agar (PDA) plates containing ampicillin (60µg/mL) and Kanamycin (50µg/mL) and incubated at 28°C in the dark. After 36 h, the fungal mycelium emerging from the xylem vessels (preferentially from the most apical stem piece) was cut into small pieces, transferred to liquid PDB and grown for additional 36 h at 28°C with orbital shaking at 170 rpm. Microconidia were collected by filtration through a nylon filter (monodur, pore diameter 10-15 µm) and 10 min centrifugation at 8243g, counted in a hemacytometer and used for dip-inoculation of tomato roots in the subsequent passage. To estimate the population bottleneck during *in planta* passaging, co-inoculations of the *Fol*4287 isolate with a hygromycin-resistant derivative were performed at different microconidial ratios (1:1, 1:0.5, 1:0.1, 1:0.05, 1:0.01 of wt versus HygR, respectively). Recovery from the xylem was performed as described above, and the presence/absence of the HygR strain was checked by plating the recovered colonies on PDA with hygromycin B.

For plate passaging, a 10 µl drop containing 10^6^ microconidia was placed at the center of a plate containing either YPDA or MMA and incubated in the dark either at 28°C or 34°C. For plates incubated at 34°C, the pH of the YPDA or MMA was buffered to 7.4 with 6.5% (v/v) 0.1 M citric acid and 43.5% (v/v) 0.2 M Na_2_HPO_4_. After 7 days, microconidia from the entire colony were collected by adding 5 ml sterile distilled water and scraping with a sterile spatula, filtered through a monodur filter and adjusted to a concentration of 10^8^/ml by counting in a hemacytometer. A 10 µl drop containing 10^6^ microconidia was then inoculated at the centre of a fresh plate for the subsequent passage. For all environmental conditions, 5 independent replicate lines were passaged in parallel. After each passage, aliquots of the collected microconidia were stored in 30% glycerol at -80°C.

### Obtaining evolved isolates from final passaged populations

After passage 10, dilutions of microconidia from each evolved population were plated on PDA media, 10 monoconidial isolates were picked randomly and subjected to two additional rounds of monoconidial isolation. After the final isolation step, small pieces from single colonies were transferred to PDB, grown for 36 h at 28°C and 170 rpm and passed through a monodur filter. The mycelium retained in the filter was used for DNA extraction while the microconidial suspensions were stored in 30% glycerol at −80°C. The presence/absence of variants of interest in the evolved isolates was determined by Sanger sequencing or PCR/RFLP analysis as described above.

### Measurement of relative fitness of evolved isolates

The relative fitness of the evolved isolates was measured by performing pairwise competition assays against the ancestor clone (wt) under different environmental conditions. Fitness in tomato plants was measured by dip-inoculating roots of tomato plants with a mixed suspension (50:50) of an evolved isolate and the ancestor clone (2.5 x 10^6^ microconidia/ml per isolate). Plants were maintained in a growth chamber as described above. After 14 days, stem pieces were cut, rinsed is sterile water and total genomic DNA was isolated using the CTAB method. The frequency of DNA from each isolate was determined by PCR with specific primers detecting differential SNVs, INDELs or TIVs as described above (Supplementary Tables 1 and 2). PCR products were separated in 1% agarose gels, imaged, quantified with the Multi Gauge 3.0 software (Fujifilm) and multiplied with the correction coefficient obtained from the corresponding standard curves. Relative fitness (*v*) of the evolved isolates *in vitro* or *in planta* was calculated using the formula *v = x2 (1- x1) / x1 (1- x2)*, where *x1* and *x2* are the frequencies of the evolved isolate before and after the competition experiment, respectively^2^. The estimated detection limit for each isolate under all environmental conditions was 0.01. Data are provided in Supplementary Table 4.

### Phenotyping of evolved isolates

For measurement of colony growth rate and conidiation, 10^6^ microconidia of each isolate were spot-inoculated on YPDA or MMA plates and incubated at 28°C in the dark. After 48 and 72 h, plates were imaged and the colony area was measured using the Multi-Gauge V 3.0 software. Growth rate was calculated as the area at 72 h minus that at 48 h. After 7 days, total microconidia were collected from the colony, filtered as described above, and quantified by counting dilutions in a hemocytometer. Cellophane invasion assays on plates were performed as described^1^. Briefly, an autoclaved cellophane membrane (colorless; Manipulados Margok, Zizurkil, Spain) was placed on top of a plate containing MMA buffered to pH 5.0 or 7.0 with 100 mM 2-(N-morpholino) ethanesulfonic acid (MES), and 10^6^ microconidia of each isolate were spot-inoculated at the centre of the plate. After 3 days incubation at 28°C in the dark, plates were imaged (before), then the cellophane membrane with the fungal colony was carefully removed, plates were incubated for 2 additional days and imaged again (after). For calculation of the invasive growth index, the area of the fungal colony after removing the cellophane was measured using the Multi-Gauge V 3.0 software. The invasive growth index was calculated using the formula (X5+X7) / (WT5+WT7), where X and WT are the growth areas after removing the cellophane in mm^2^, of the evolved isolate or the wt strain, respectively, at pH 5 and pH7, respectively. All plate phenotyping experiments were performed at least two times, each with two replicate plates.

For plant infection assays, roots of 2-week-old tomato plants (cultivar Monika) were dip-inoculated using an inoculum concentration of 5 x 10^5^ microconidia/ml, or sterile water as a control, and incubated in a growth chamber as described above. Groups of 10 plants were used for each treatment. The number of dead individuals was recorded daily for at least 35 days. Infection of *Galleria mellonella* larvae was done as described^3^ by injecting 8 μL of a microconidial suspension containing 2 x 10^6^ microconidia/ml, or PBS as a control, into the hemocoel through the left proleg, using a Burkard Auto Microapplicator (Burkard Manufactoring Co. Limited, Hertfordshire, UK) with a 1 ml syringe. Groups of 20 larvae were used for each treatment. Larvae were incubated in glass containers at 30°C and the number of dead individuals was scored daily. For both types of infection assays, survival was calculated using the Kaplan–Meier method and compared among groups using the log-rank test. Data were analyzed with the software GraphPad Prism 4. The mortality index was calculated using the formula (AC-AX) / (AC-AWT) where AC is the area under curve (AUC) of the water control, and AX and AWT are the AUC of the plants inoculated with the evolved isolate or the wt strain, respectively. All infection experiments were performed at least two times. Phenotyping data are provided in Supplementary Table 4.

### Quantification of conidial size and cell density-dependent inhibition of germination

The relative size of microconidia of the wild type and the *21009*Δ strain was measured by flow cytometry using a BD FACSCelesta™ with BD FACSDiva™ Software, version 9.0 (Becton Dickinson Biosciences, Franklin Lakes, NJ, United States), following a protocol described^4^. Briefly, 1.5×10⁶ fresh microconidia from each strain were resuspended in 1 ml of 50 mM sodium citrate, pH 7.1. To ensure the accuracy of the measurements, two successive gatings were applied to consider only viable single-cell events for data acquisition. First, cell viability staining was performed by adding 20 µM propidium iodide directly to the sample followed by 5 min incubation in the dark. Second, cell multiplets were identified and excluded from the final dataset. A total of 100,000 events were acquired for each sample. Channel exported linear-scaled forward scatter data were used to determine microconidia size with FlowJo flow cytometry analysis software version 10 (Tree Star, Ashland, OR, USA). Images of microconidia samples were obtained using the AxioVision 4.8 software and a Zeiss Axio Imager M2 microscope (Zeiss, Oberkochen, Germany) equipped with an Evolve Photometrics EM512 digital camera (Photometrics Technology, Tucson, AZ, United States).

Inhibition of conidial germination at high-cell densities was measured as described ^5^. Briefly, fresh microconidia were inoculated in germination medium (GM) either at optimum (3.19 x 10^6^ ml^-1^) or inhibitory concentration (8.6 x 10^7^ ml^-1^) and incubated at 28°C and 170 rpm for 13 or 15 h, respectively. To quantify germination, images were acquired in a Zeiss Axio Imager M2 microscope as described above, and the number of germinated conidia was counted. At least 200 conidia were scored per strain and experimental condition. Two independent experiments were performed, each with three replicates. Statistical analysis was performed using two-way ANOVA followed by Dunnett’s multiple comparison test (Padj < 0.05). Quantitative data are provided in Supplementary Table 5.

### DNA extraction and whole-genome sequencing of the ancestor clone and the passaged populations

Genomic DNA was extracted as described using the CTAB method^1^. The quality and quantity of the DNA was routinely determined by running aliquots in agarose gels and by spectrophotometric analysis in a NanoDrop ND-1000 spectrophotometer (NanoDrop Technologies, Wilmington, DE, USA). Library preparation and whole genome shotgun sequencing of the ancestor clone and the final passaged populations were performed at the Tufts University Core Facilities with an Illumina HiSeq 2500 Platform (2×71 cycles), except for the minimal media-passaged populations, which were sequenced at the Genomics Resource Laboratory of the University of Massachusetts (UMass) Amherst with an Illumina NextSeq 500 platform (2×75 cycles). To check whether the more error-prone NextSeq 500 platform introduced any bias to the analysis, the minimal media-passaged population M4-10 was resequenced with an Illumina MiSeq platform (2×75 cycles) at the Genomics Resource Laboratory UMass. FastQC version 0.11.5 was used to check raw read quality^6^.

### Genome assembly and mapping

For all analyses, the assembled and annotated *Fol*4287 genome was used as reference (accession QESU00000000) (Ayhan et al. 2018). The raw reads from sequenced populations and the paired-end Illumina reads of the ancestor clone *Fol*4287 (wt) were mapped to the reference genome using BWA mem (version 0.7.15) with the following options ‘-t 8 -M -a -v 1’^7^. The alignments were processed using Picard (version 2.17.8) CleanSam, Samtools view (version 1.3), Picard FixMateInformation, Picard MarkDuplicates, and Samtools index using the default options^8,9^. Samtools flagstat and Picard CollectRawWgsMetrics were used to assess mapping quality. For wt and the final passaged populations, the base quality scores were recalibrated via GATK BaseRecalibrator (version 3.57) using initial variant calling with FreeBayes (version v1.0.2-33-gdbb6160) for ‘--knownSites’ option^10,11^.

### Single nucleotide variant (SNV) and small insertion deletion (INDEL) calling

SNVs and INDELs were called by GATK Mutect2 (version 4.1.4.1) using the wt sample as “normal” and otherwise the default options. The one command line per sample group was run (e.g., the samples wt, Y1-10, Y2-10, Y3-10, Y4-10, and Y5-10 were analyzed together). The calls were filtered via GATK FilterMutectCalls using the default options. All detected variants were manually inspected and further filtered using Integrative Genomics Viewer (IGV) ^12^. The allele frequency (AF) values for high coverage regions were adjusted to reflect real frequencies. For each detected variant in the sequenced samples of the intermediate populations, allele frequency for each population was reported even if it was not detected by these tools. AFs lower than 0.1 were filtered out for most of the analyses.

### TIV calling

TE insertions were detected with TEfinder^13^. TE annotations were performed *de novo* and classified using RepeatModeller version 1.0.11^14^ and a curated TE library^15^. RepeatMasker version 4.0.7 was used to mask and annotate the genome sequence. TIVs were detected by TEfinder and further filtered out by testing their presence in wt using BEDtools intersect with the option ‘-v’ and by manual inspection with Integrative Genomics Viewer^16^. TIV allele frequencies were estimated manually.

### CNV calling

BEDtools genomecov with the option ‘-d’ was used to output the read depth at single nucleotide resolution for the final populations and wt. Then, a custom MatLab code was used to calculate median coverages of 10 kb windows of the genome. Read coverages were normalized to the median read coverages of the samples and the final population samples were normalized to wt. Hierarchical clustering was performed with the normalized copy numbers of the loci showing >20% change in any sample, and the AR contigs and some unmapped contigs were ordered accordingly. This ordering was used for both read coverage and genomic feature maps.

The coverages of the small arm of chromosome 13, of chromosome 15, as well as of the duplicated region of chromosome 3 were multiplied by 2 to reflect the real copy numbers. Regions with >10% changes were considered CNVs. For the circular map, normalized read depths in 10 kb windows of all final populations were averaged. Total genome sizes were estimated from the raw 10 kb windows normalized by sample median coverage (Estimated genome size: 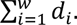 10000, where *i* is the 10 kb windows, *w* is the total number of 10 kb windows, and *d_i_* is the median read depth of the 10 kb window normalized to overall median read coverage). One-way ANOVA was performed in three passaging groups on MatLab.

### Genomic Features

For analysis of TE content, simple repeats were removed from the previously identified repeats. BEDtools genomecov with the option ‘-d’ and a custom MatLab code were used to calculate the median TE content over 10 kb windows. Mean GC content of the genome was calculated over 10 kb windows using a MatLab code.

ChIP-seq raw reads for H3K4me2 and H3K27me3 in *Fol*4287 were retrieved from NCBI GEO accession GSE1212839^17^. The reads were mapped to the *Fol*4287 genome using BWA mem and the alignments were processed as previously described. The read coverage was detected using BEDtools genomecov with the option ‘-d’. The reads from the two experimental replicates were added and genome coverage was calculated as the median values over 1 kb windows.

One-tail Fisher’s exact test was used to test the null hypothesis that there are no non-random associations between the variants (TIVs or SNVs & INDELs) and sites with H3K27me3. Circos version 0.69-6 was used to plot the circular genome map^18^.

### TE sequence alignment

Sequence alignment of *Hornet* and *Hormin* TEs was performed using NCBI Blast^19^. The domains were annotated by aligning them to a Restless-like transposase (Uniprot accession: A0A395M6G3) using NCBI Blast and Conserved Domain-search^20^.

### Independent confirmation and frequency quantification of variants by PCR

The presence of selected SNVs and INDELs in the passaged lines was independently confirmed by PCR amplification of genomic DNA using specific primers flanking the site of each event (Supplementary Table 1), followed by Sanger sequencing of the amplified fragment or, when possible, by restriction fragment length polymorphism (RFLP) analysis in agarose gels. Larger INDELs and TIVs were confirmed by PCR amplification of genomic DNA using specific primers flanking the site of the event (Supplementary Table 1), followed by electrophoresis in agarose gels and determination of the fragment size. In all cases, PCR reactions were performed using the Expand High Fidelity PCR System (Roche Life Sciences). Reactions contained 300 nM of each primer, 2.5 mM MgCl2, 0.8 mM dNTP mix, 0.05 U/μl polymerase and 5-10 ng/μl genomic DNA. PCR cycling conditions were as follows: an initial step of denaturation (5 minutes, 94°C); 35 cycles of 35 seconds at 94°C, 35 seconds at the calculated primer annealing temperature and 1 minute/1.5 Kb extension at 72°C (or 68°C for templates larger than 3 kb); and a final extension step of 10 minutes at 72°C (or 68°C). For RFLP analysis, PCR fragments were treated for 2-4 hours with 2 units per 100 ng DNA of the appropriate restriction enzyme using the conditions recommended by the manufacturer. Negative controls (uncut PCR bands or no DNA template) were included. To estimate the allele frequencies of SNVs, INDELs and TIVs in a population, the obtained DNA fragments were separated in 1% agarose gels, imaged and quantified using the Multi Gauge 3.0 software (Fujifilm). Where INDEL and TIV variants produced PCR fragments of different sizes, a standard curve was elaborated using known DNA mixtures containing the proportions of the wt and the variant allele of 0:100, 10:90, 20:80, 30:70, 40:60, 50:50, 60: 40, 70:30, 80:20, 90:10 and 100:0. DNA quantification for standard curves was carried out fluorimetrically using the Quant-iT DNA Assay Kit Broad Range kit (Molecular Probes Inc., Leiden, The Netherlands) in a Tecan Safire fluorospectrophotometer (Tecan España, Barcelona, Spain). Finally, frequencies obtained by image analysis were multiplied with the correction coefficient obtained from the corresponding standard curve.

Frequencies of CNVs in a population were estimated by quantitative real time PCR (RT-qPCR) using a CFX Connect^TM^ Real-Time PCR System (BioRad) with the FastStart Universal SYBR Green Master kit (Roche). Copy numbers were calculated using the double delta Ct method^21,22^, using the single copy gene *FOXG_01569* (actin) as a reference, and expressed relative to the ancestor clone. All RT–qPCR experiments were performed with three independent biological repeats, each of them with three technical repeats. Primers used are listed in Supplementary Table 1.

### RNA-sequencing for measuring TE transcript levels

100 ml PDB, BD Difco, USA) was inoculated with 20 μl frozen stocks of *Fol*4287 and incubated at 28°C, 140 rpm for 4 days. The mycelium was harvested in a paper filter, rinsed with sterile water, and dried using sterile paper towels. 200 mg mycelium was mixed with 20 ml YPD, pH 5.0 or 7.4 as described above in 50 ml tubes, incubated 60 min at 28°C and 34°C, respectively, and filtered and dried as described above. For 34°C treatments, the filtration process was performed in the incubator.

100 mg mycelium was homogenized in 1 ml TRIzol Reagent (Invitrogen, USA) using the 1.5 ml Next Advance Pink RINO RNAse-free lysis kit and homogenized in a Next Advance Bullet Blender Tissue Homogenizer at speed 10. Samples were incubated in ice for 1 min and the homogenizing step was repeated, followed by 5 min incubation at room temperature. Then, 0.2 ml chloroform was added to the samples and mixed. After 2–3 min, samples were centrifuged 18 min at 10,000×g and 4°C. The top phase was transferred to a new 1.5 ml RNase-free tube. 500 ml of ethanol was added. Samples were loaded on RNeasy columns (RNeasy Mini Kit, QIAGEN) following the manufaturer’s instructions and the RNA was eluted in 30 μl of RNase-free water and stored at - 20°C. The RNA was quantified using Qubit RNA BR Assay and the quality was checked with an Agilent Bioanalyzer. The samples were sequenced at the Genomic Resource Laboratory, UMass on an Illumina NextSeq 500 platform with 2×76 cycles.

The reads were mapped to the *Fol*4287 genome sequence using Hisat2 (version 2.0.5) with the options ‘-p 15 -X 500 --dta’^23^. Alignments were sorted with Samtools (version 1.3) ^24^ sort, duplicates were marked with Picard (version 2.0.1) MarkDuplicates and indexed by Samtool index. TEcount (version 2.2.1) was used with default options to generate the read counts mapped to TEs. Finally, the read counts were normalized across samples by calculating the geometric means across sample sets and normalizing each sample by their median in a custom MatLab code.

### Quantification of transposase gene expression by reverse transcriptase RT-qPCR analysis

To measure transcript levels of the *Hornet* transposase gene, total RNA was isolated from snap frozen tissue and used for reverse transcriptase quantitative PCR (RT-qPCR) analysis as described^1^. Briefly, RNA was extracted using the Tripure Reagent and treated with DNase (both from Roche). The resulting RNA was reverse transcribed with the iScript^TM^ cDNA Synthesis Kit (Bio-Rad) to synthesize the cDNA, and qPCR was carried out using the FastStart Essential DNA Green Master (Roche) in a CFX Connect Real-Time System (Bio-Rad) according to the manufactureŕs instructions. Data were analyzed using the double delta Ct method by calculating the relative transcript level normalized to the *act1* gene (*FOXG_01569*). Three biological replicates with three technical replicates each were conducted for each experimental condition. Primers used and transcript level data are listed in Supplementary Table 1.

### Data and code availability

All quantitative data and statistical analyses are provided in Supplementary Tables 1–5. The genome sequencing reads are deposited in NCBI SRA (Accession: PRJNA682786). Bash scripts and the MatLab codes are available at https://github.com/d-ayhan/Foxysporum_STEE.

## Acknowledgements

We are grateful to Drs. Toni Gabaldón and Elena Pérez Nadales for initial discussions leading to the passage and re-sequence experiments. María Ortega and Mariló Alcaide Caballero are acknowledged for technical assistance. CLD was supported by Ph.D. fellowship FPI EEBB-I-16-BES-2014-070450 from MICINN and by a postdoctoral contract from Junta de Andalucía (PAIDI 2020). ADL and LGG were supported by Ph.D. fellowships FPU19/04366 and FPU16/01029, respectively, from the Spanish Ministry of Universities. This work was funded by grants PID2019-108045RB-I00, PLEC2021-007777, TED2021-130262B-I00, PDC2022-133749-I00 and PID2022-140187OB-I00 from the Spanish Ministry of Science and Innovation (MICINN) to ADP; and by grants EY 030150 from the United States National Institute of Health National Eye Institute, 1652641 from the United States National Science Foundation, and MAS00612 from the United States Department of Agriculture National Institute of Food and Agriculture to LJM.

## Author Contributions

**Conceptualization:** CLD, DHA, LJM, ADP. **Data curation:** CLD, DHA, ARL, LGG, LJM, ADP. **Formal analysis:** CLD, DHA, ARL, LGG, LJM, ADP. **Funding acquisition:** LJM, ADP. **Investigation:** CLD, DHA, ARL, LGG, LJM, ADP. **Methodology:** CLD, DHA, ARL, LGG, LJM, ADP. **Project administration:** LJM, ADP. **Resources:** LJM, ADP. **Software:** DHA, LJM. **Supervision:** CLD, DHA, LJM, ADP. **Validation:** CLD, DHA, ARL, LGG, LJM, ADP. **Visualization:** CLD, DHA, ARL, LGG, LJM, ADP. **Writing – original draft preparation:** CLD, DHA, LJM, ADP. **Writing – review and editing:** CLD, DHA, ARL, LGG, LJM, ADP.

## Supplementary Information

**Supplementary Table 1.** Various data.

**Supplementary Table 2.** List of variants.

**Supplementary Table 3.** Relative Fitness calculations

**Supplementary Table 4.** Phenotyping of evolved isolates

**Supplementary Table 5.** Phenotyping of 21009delta mutant

## Extended Data Figure legends

**Extended Data Figure 1.**
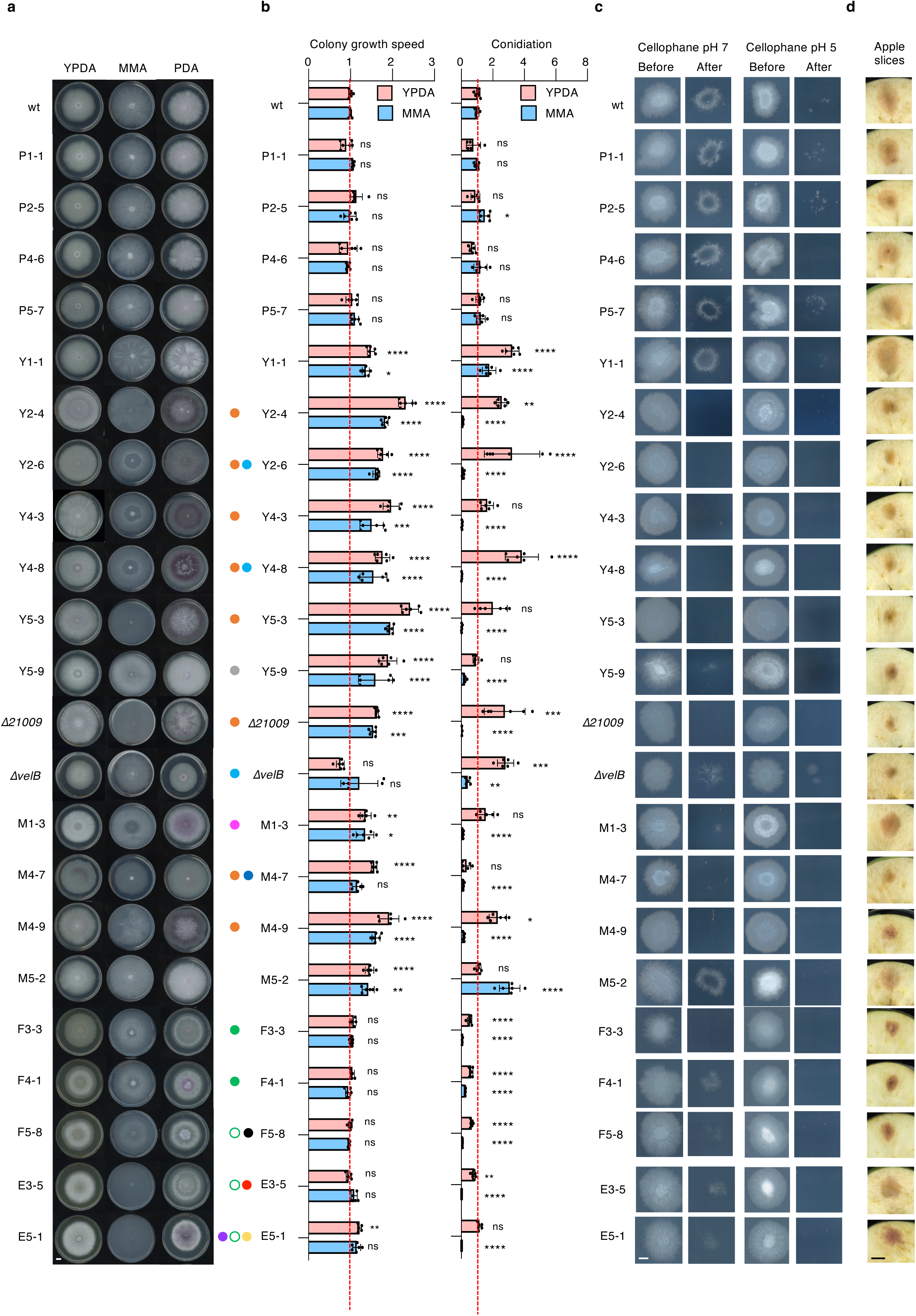
Adaptation to plates results in increased proliferation at the cost of reduced invasive growth. **a, b.** Colony phenotypes on different media (a); and growth speed and conidiation (b) of the indicated monosporic isolates obtained from populations submitted to ten serial passages through tomato plants (P), YPDA (Y,F) or MMA plates (M,E) either at 28°C (Y,M) or 34°C (F,E). Each row represents an individual isolate. Isolates were spot-inoculated on the indicated media and grown for 7 days at 28°C. Growth speed was calculated as the difference between the colony area on day 2 and 3. Values in (b) are normalized against the mean of the ancestor clone (wt). Error bars, SD. Results shown are from 2 independent experiments with 3 replicates each. *P < 0.05, **P < 0.01, ***P < 0.001, ****P < 0.0001, versus wt according to Bonferroni’s multiple comparisons test. **c, d.** Invasive growth through cellophane membranes (c) or on apple slices (d). Isolates were spot-inoculated on PDA plates buffered pH 5 or 7 with 100 mM MES and covered with a cellophane membrane, grown for 2 days at 28°C (before), then the membrane with the fungal colony was removed and plates were incubated for an additional day to visualize the presence of fungal mycelium that had penetrated through the cellophane (after) (c); or on apple slices and incubated 3 days at 28°C and 100% humidity (d). Images shown are representative of 2 independent experiments with 3 replicates each. Scale bars, 1 cm. Colored dots next to isolate numbers indicate mutations in genes mutated in more than one independent passaged population (see Table 1 for the list and color legend of recurrently mutated genes). Empty and solid green dots represent the TIVs in gene *FOXG_21943* in hetero- and hemizygosis, respectively.

**Extended Data Figure 2.**
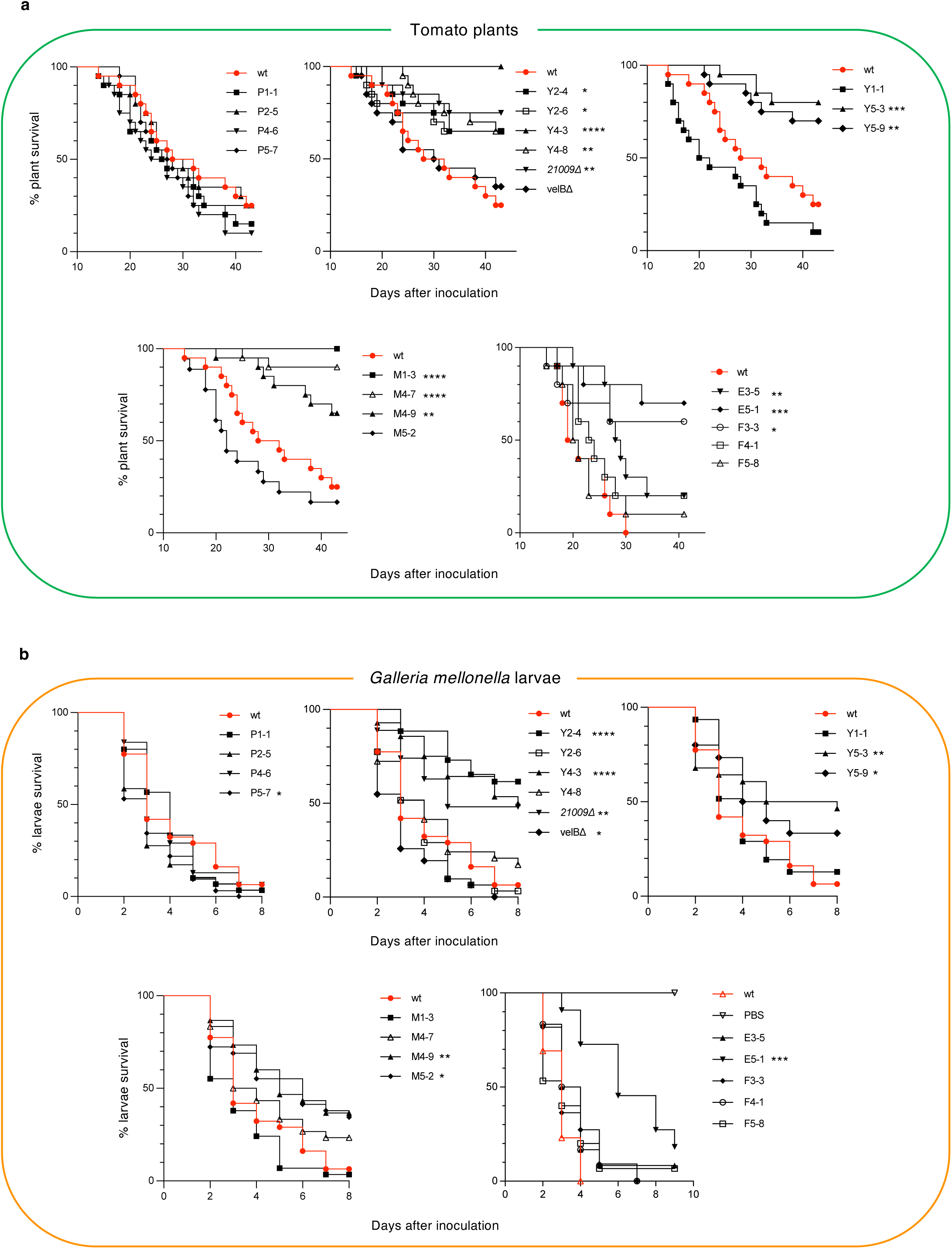
Adaptation of *Fol4287* to plates results in reduced virulence on plant and animal hosts. **a, b.** Kaplan-Meier plots showing survival of tomato plants (a) or *Galleria mellonella* larvae (b) inoculated with the wt (red) or the indicated isolates from populations passaged through tomato plants (P) or through YPDA or MMA plates at 28°C (Y or M, respectively) or at 34°C (F or E, respectively); or with the *FOXG_21009* or *velB* knockout mutants. n = 20 plants or 30 larvae per group. *P < 0.05, **P < 0.01, ***P < 0.001, ****P < 0.0001 versus wt according to Log-Rank test. Infection experiments wre performd at least twice with similar results.

**Extended Data Figure 3.**
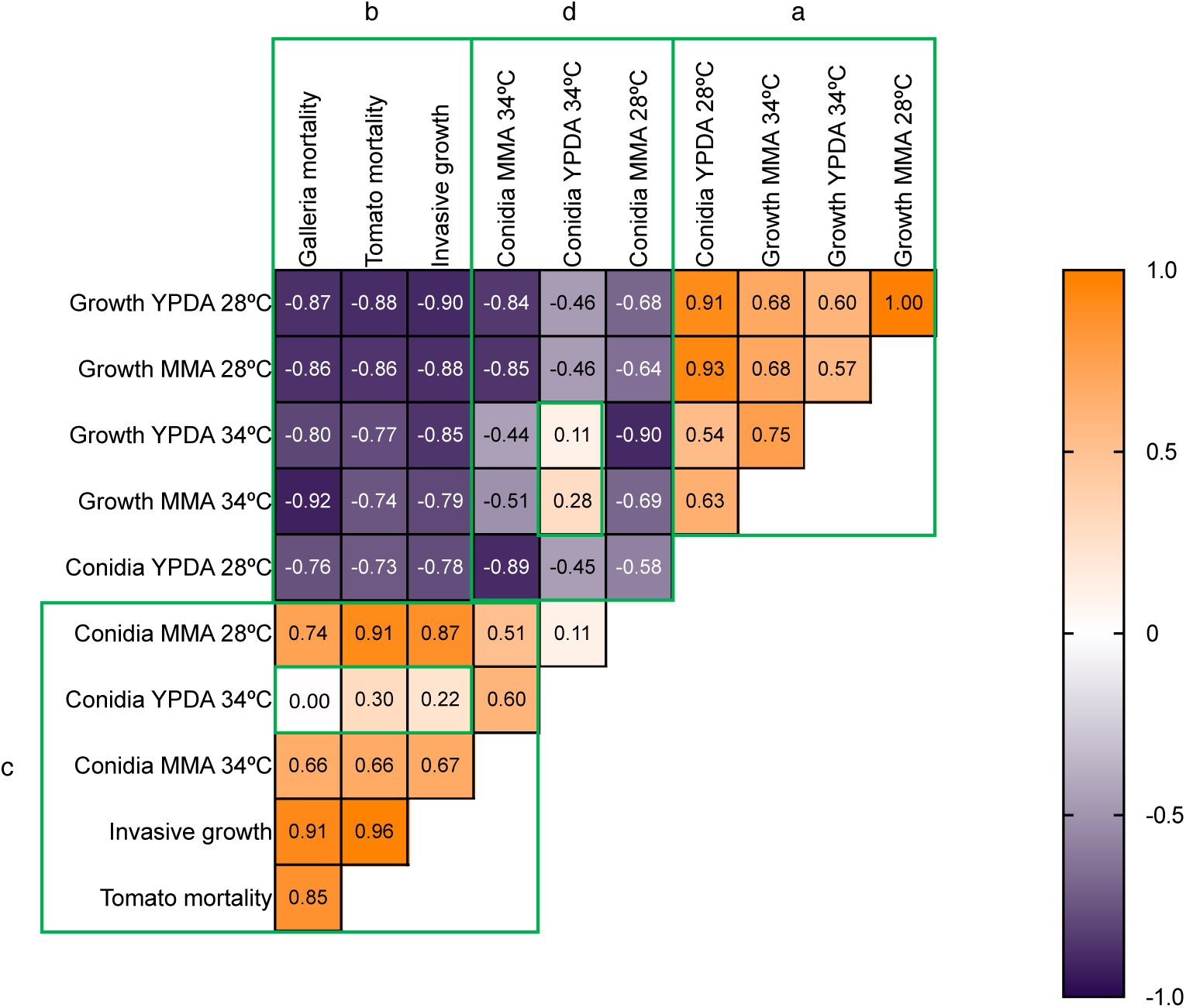
Correlation between different phenotypes in evolved isolates. **a.** Pearson r correlation matrix between the indicated phenotypes measured in the evolved isolates (see Fig. 1a and Extended Data Fig. 1). Calculations were performed after first computing the mean of side-by-side replicates, and then analyzing those means. n = 20. Green boxes highlight groups of phenotypes with similar correlations.

**Extended Data Figure 4.**
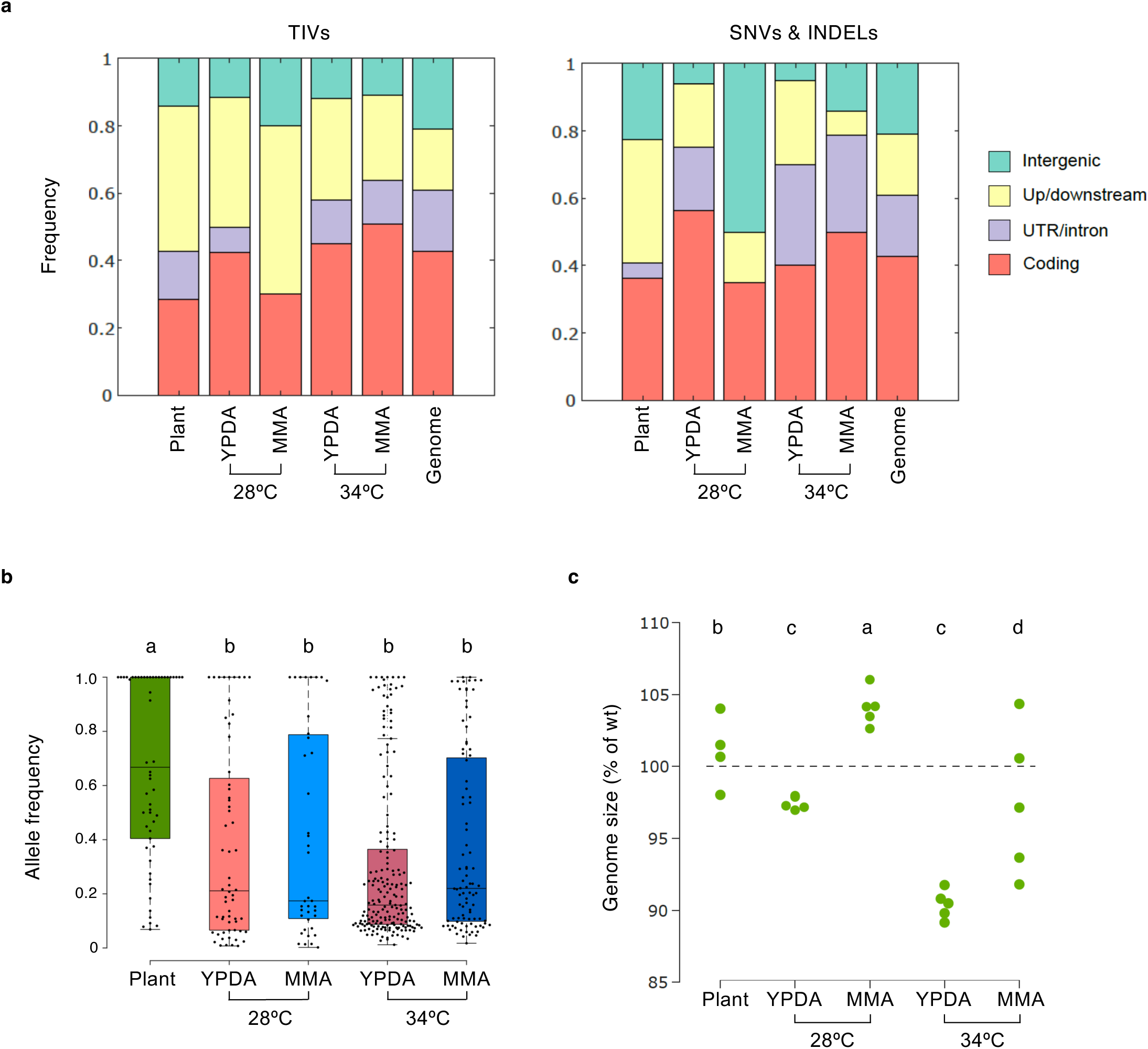
Distribution and dynamics of mutations and genomes in evolved populations. **a.** Relative distribution of TIVs (left) and SNVs/INDELs (right) in populations submitted to ten serial passages through the indicated conditions, to coding sequences (CDS), UTR or intronic regions, regions encompassing 1 kb up- or downstream of genes, or intergenic regions. Only mutations with allele frequency (AF) ≥ 0.1 were considered. **b.** AF distribution of mutations in the evolved populations submitted to ten serial passages through the indicated conditions. Only mutations with AF ≥ 0.1 are shown. AF distributions with the same letter are not significantly different according to One-way ANOVA (p<0.0001). **c**. Changes in genome size of the evolved populations submitted to ten serial passages through the indicated conditions. Genome sizes are normalized to that of the ancestor clone (wt, dotted line). Each dot represents an evolved population. Groups with the same letter are not significantly different according to One-way ANOVA (*P* = 9.8 x 10^-5^).

**Extended Data Figure 5.**
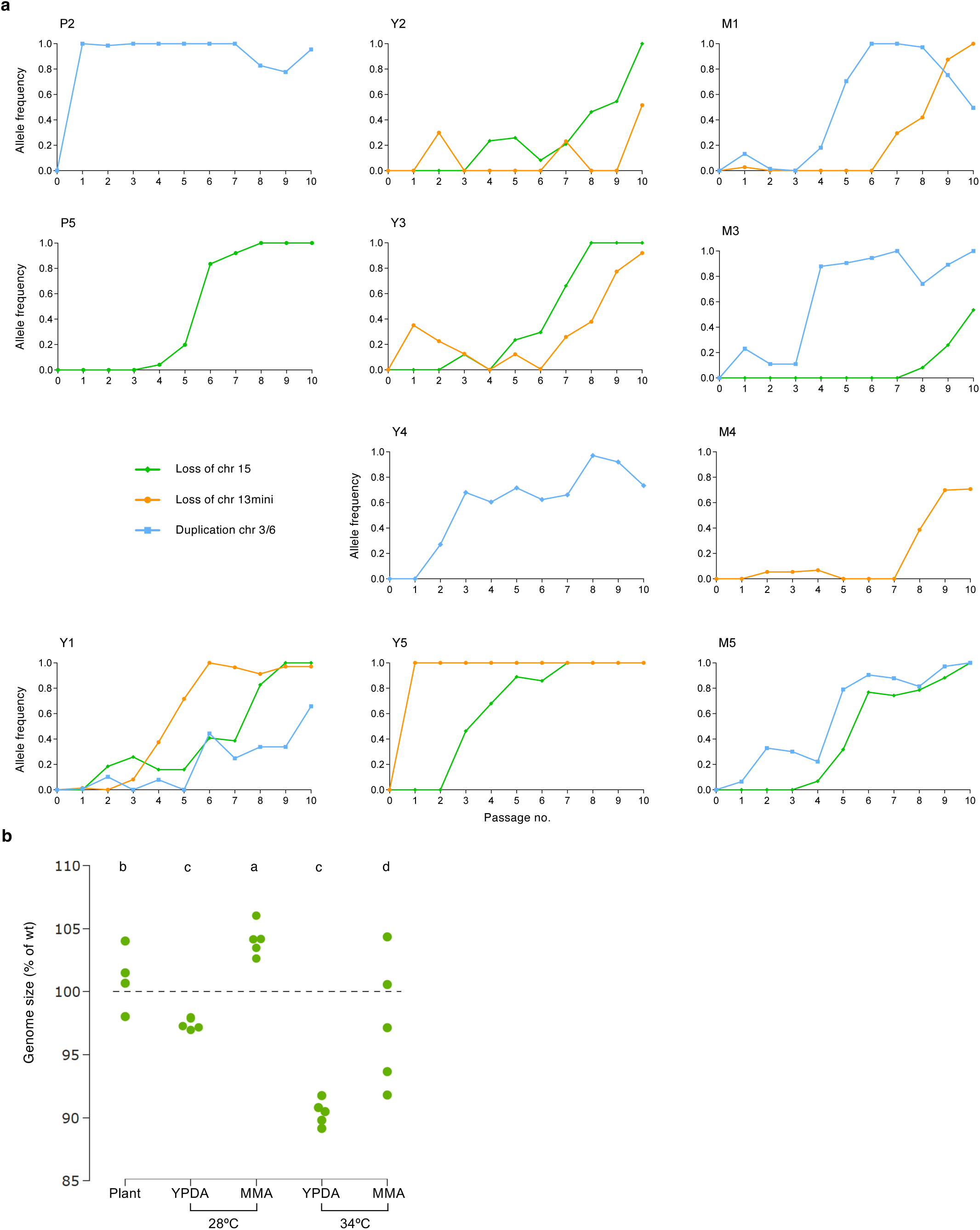
Evolutionary dynamics of recurrent CNVs detected in experimentally evolved populations and their effect on genome size. Frequency dynamics of the indicated CNVs in independently evolved populations submitted to ten consecutive passages on tomato plants (P), YPDA (Y) or MMA plates (M) at 28°C. Loss of chr 15 or chr 13mini refers to loss one copy of a duplicated accessory region coinciding with chromosome 15 or 13mini, respectively; Duplication chr 3/6 refers to duplication of a region located on accessory chromosome 3 to generate the new accessory chromosome 6. Ploidy levels of the relevant genomic regions were measured by real time qPCR of genomic DNA, calculated using the ΔΔCt method and normalized to the single copy actin gene. Note that some populations carry multiple CNVs. **b.** Changes in genome size of the evolved populations after ten serial passages through tomato plants, or YPDA or MMA plates, either at 28°C or 34°C. Genome sizes are normalized to that of the ancestral clone (wt, dotted line). Each dot represents an evolved population. Groups with the same letter are not significantly different according to One-way ANOVA (*P* = 9.8 x 10^-5^).

**Extended Data Figure 6.**
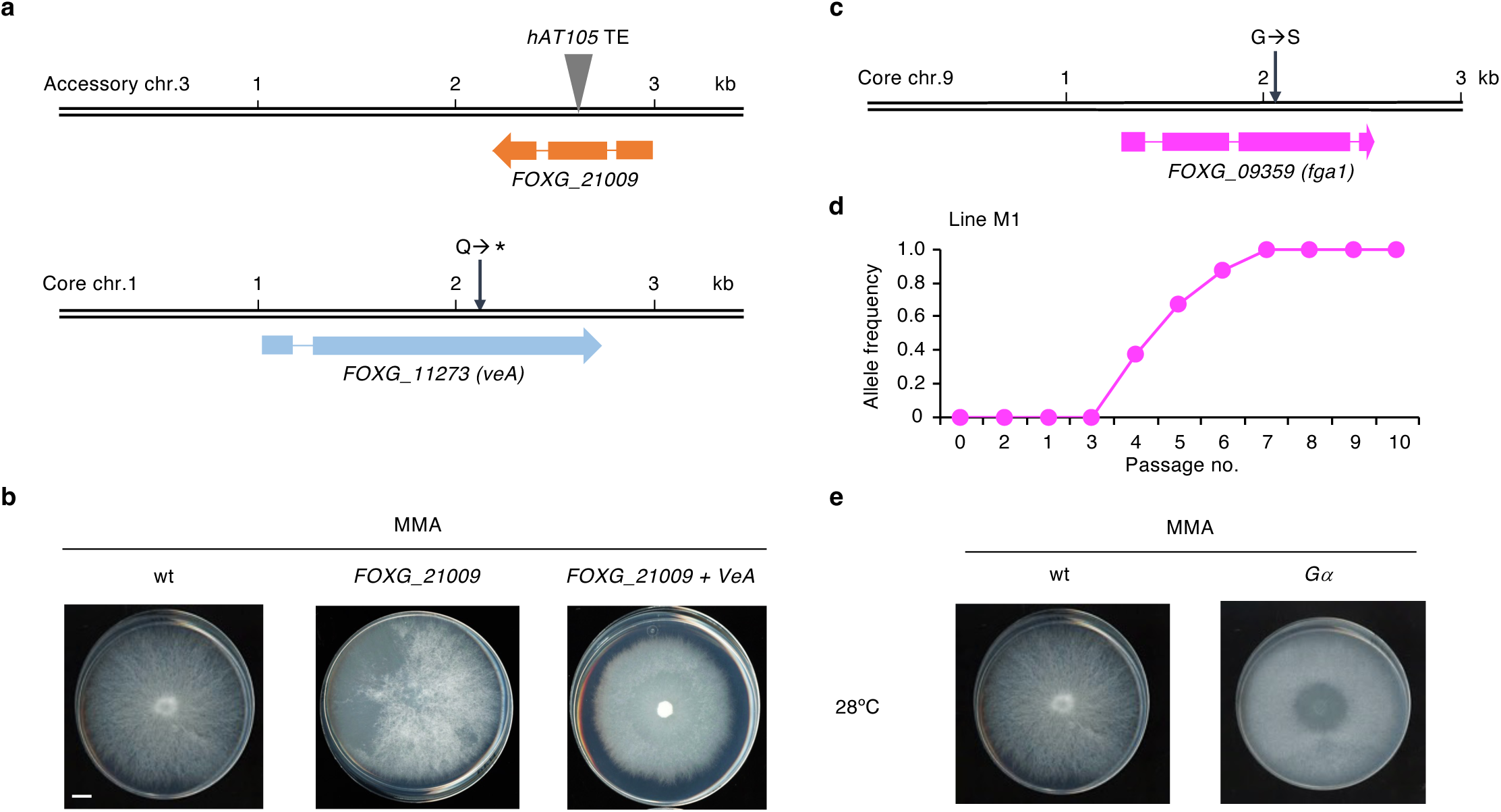
Adaptive mutations selected during serial passaging on plates lead to altered colony morphology. **a,c.** Diagrams depicting adaptive mutations selected in populations passaged on MMA at 28°C. (a, upper) Insertion of the TE *hAT105* in gene *FOXG_21009* located on accessory chromosome 3, encoding a putative iron 2-oxoglutarate-dependent dioxygenase. (a, lower) Nonsense mutation in gene *FOXG_11273* located on core chromosome 1, encoding the VeA subunit of the Velvet regulatory complex. (b) Missense mutation in gene *FOXG_09359* located on core chromosome 9, encoding the Ga subunit of heterotrimeric G proteins Fga1, causing a dominant active version of the protein. **b,e.** Representative images of colony phenotypes on MMA of the ancestor clone (wt) and isolates from plate-passaged populations that carry the mutations depicted in (a) and (c), respectively. Scale bar, 1 cm. **d.** Allele frequency dynamics of the *FOXG_09359* mutation in the MMA-passaged population M1. **f.** Diagram depicting five independent insertion events of the TE *Hormin* in gene *FOXG_21943* encoding a putative prolyl hydroxylase, selected in populations passaged on YPDA or MMA at 34°C, pH 7.4. Note that all TIVs map close to the 3’-ends of the *FOXG_21943* transcript and the *FOXG_15014* CDS. Both genes are present in two copies on ARs. **g.** Representative images of colony phenotypes on YPDA or MMA at 34°C, pH 7.4 of the ancestor clone (wt) and two isolates from YPDA or MMA plate-passaged populations, respectively, each carrying one of the mutations depicted in (f).

**Extended Data Figure 7.**
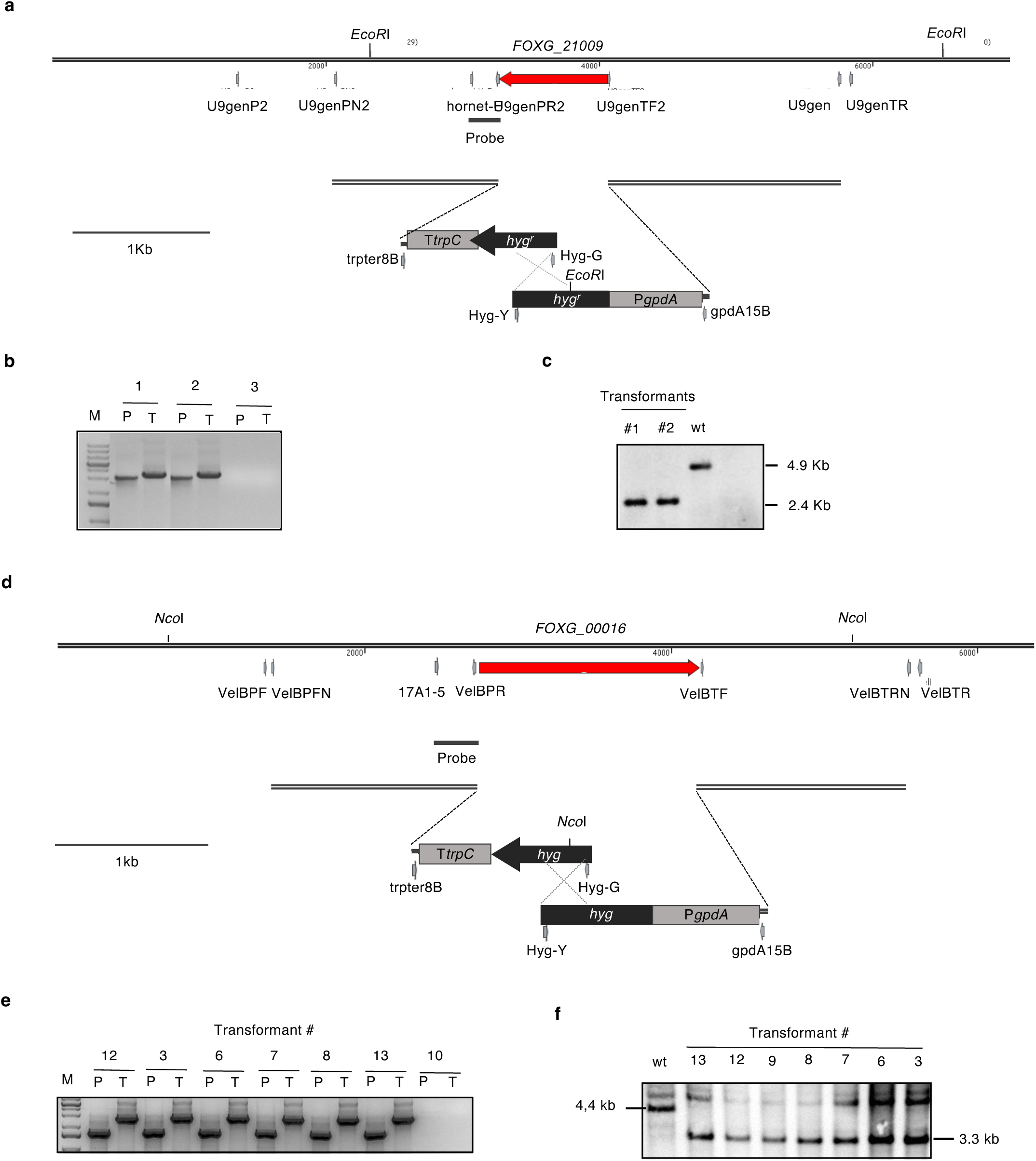
Targeted replacement of the *FOXG_21009* and the *velB* genes. **a.** Physical maps of the *F. oxysporum FOXG_21009* locus and of the split-marker gene replacement constructs obtained by fusion PCR. Relative positions of restriction sites and PCR primers are indicated. *hyg*, hygromycin resistance gene; *PgpdA,TtrpC*, gpdA promoter and trpC terminator, both from *Aspergillus nidulans*. **b.** PCR amplification of genomic DNA of independent transformants using primer pairs VelBPF/Hyg-G (P) or trpter8B/VelBTR (T) indicated in (a). Presence of amplification bands indicates homologous insertion of the deletion construct. **c.** Southern blot analysis of genomic DNA of the wild type strain (wt) and seven independent transformants, treated with *Nco*I, separated on a 0.7% agarose gel, transferred to a nylon membrane and hybridized with the DNA probe indicated in (a). Molecular sizes of hybridizing bands are indicated. **d.** Physical maps of the *F. oxysporum velB* (*FOXG_00016*) locus and of the split-marker gene replacement constructs obtained by fusion PCR. Relative positions of restriction sites and PCR primers are indicated. *hyg*, hygromycin resistance gene; *PgpdA,TtrpC*, gpdA promoter and trpC terminator, both from *Aspergillus nidulans*. **e.** PCR amplification of genomic DNA of independent transformants using primer pairs VelBPF/Hyg-G (P) or trpter8B/VelBTR (T) indicated in (a). Presence of amplification bands indicates homologous insertion of the deletion construct. **f.** Southern blot analysis of genomic DNA of the wild type strain (wt) and seven independent transformants, treated with *Nco*I, separated on a 0.7% agarose gel, transferred to a nylon membrane and hybridized with the DNA probe indicated in (a). Molecular sizes of hybridizing bands are indicated.

## Notes

### Competing Interest Statement

The authors have declared no competing interest.

